# Modular vector assembly enables rapid assessment of emerging CRISPR technologies

**DOI:** 10.1101/2023.10.25.564061

**Authors:** Abby V. McGee, Yanjing V. Liu, Audrey L. Griffith, Zsofia M. Szegletes, Bronte Wen, Carolyn Kraus, Nathan W. Miller, Ryan J. Steger, Berta Escude Velasco, Justin A. Bosch, Jonathan D. Zirin, Raghuvir Viswanatha, Erik J. Sontheimer, Amy Goodale, Matthew A. Greene, Thomas M. Green, John G. Doench

## Abstract

The diversity of CRISPR systems, coupled with scientific ingenuity, has led to an explosion of applications; however, to test newly-described innovations in their model systems, researchers typically embark on cumbersome, one-off cloning projects to generate custom reagents that are optimized for their biological questions. Here, we leverage Golden Gate cloning to create the Fragmid toolkit, a modular set of CRISPR cassettes and delivery technologies, along with a web portal, resulting in a combinatorial platform that enables scalable vector assembly within days. We further demonstrate that multiple CRISPR technologies can be assessed in parallel in a pooled screening format using this resource, enabling the rapid optimization of both novel technologies and cellular models. These results establish Fragmid as a robust system for the rapid design of CRISPR vectors, and we anticipate that this assembly approach will be broadly useful for systematic development, comparison, and dissemination of CRISPR technologies.

## INTRODUCTION

CRISPR technology is routinely used for a diversity of genome editing applications^1^. In addition to directing site-specific dsDNA breaks, tethering functional domains to Cas proteins enables regulating gene expression via transcriptional activation and interference; modifying chromatin marks; and editing DNA directly via base editing and prime editing^2,3^. Cas enzymes from diverse evolutionary lineages^4^ have been developed for genome engineering, including Cas9 enzymes that differ in size and PAM requirements; other Cas proteins that have novel activities, such as Cas12a, which has the ability to self-process arrays of guides; and enzyme families that target RNA rather than DNA^5^. Further still, the naturally-occurring enzymes have been rationally engineered to alter their properties, such as expanding the PAM sequence or decreasing tolerance for off-target activity^6^. Finally, these technologies can be deployed at scale to systematically perturb gene function^7,8^.

This explosive increase in experimental options can generate decision paralysis for researchers looking to implement these tools in service of their biological questions. Further, practical barriers may limit the ability to test a newly-described technology in a model system different from that in which it was first developed, as the components are often not in the desired expression architecture, causing researchers to either modify an existing vector via custom cloning or compromise their preferred experimental plan to fit an existing vector. PCR-based cloning is generally unique to each new cloning campaign, requiring the use of custom primers that are rarely repurposable. In contrast, Golden Gate (GG) cloning, which relies on Type IIS restriction enzymes with programmable overhangs, lends itself to modular systems using interchangeable and reusable parts^9–11^. A robust CRISPR toolkit has been developed for plants^12^, and another focused on engineering synthetic transcriptional circuits in mammalian systems included CRISPR elements^13^, yet there remains an unmet need for rapidly deploying the latest in CRISPR technology in mammalian cells.

Here we develop Fragmid, a toolkit of fewer than 200 modular fragments that can be mixed and matched to create millions of possible vectors for a variety of CRISPR applications, including knockout, activation (CRISPRa), interference (CRISPRi), base editing, and prime editing. This system is compatible with many delivery mechanisms, including lentivirus, PiggyBac transposon, and AAV. Modular design allows for the rapid addition of new fragments as novel enzymes and applications are published. We demonstrate the utility of Fragmid by comparing lentiviral destination vectors, performing head-to-head Cas9 ortholog comparisons via AAV delivery, and developing a pooled screening approach for high-throughput assessment of CRISPR technologies. Finally, we accompany this resource with a web portal to enable users to browse existing components and rapidly iterate on vector design.

## RESULTS

The Fragmid toolkit is comprised of destination vectors into which modular fragments are assembled. Destination vectors contain an origin of replication and a resistance marker for propagation in bacteria, as well as features necessary for the delivery and expression of inserted fragments in target cells, such as lentiviral elements for subsequent viral packaging and transduction of mammalian cells. Fragment vectors fall into six main categories of components necessary for CRISPR technology: Guide cassettes, RNA polymerase II (Pol II) promoters, N’ terminus domains, Cas proteins, C’ terminus domains, and 2A-selection markers (Figure 1a). Guide cassette fragments typically include an RNA polymerase III (Pol III) promoter and a constant RNA sequence that allows for the association of a guide RNA with the appropriate Cas protein, e.g., a tracrRNA-derived sequence for Cas9 or direct repeat (DR) for Cas12a. Fragments contain either validated targeting guides to serve as positive controls or BsmBI-based cloning sites to enable subsequent cloning of individual guides or pooled libraries. Pol II promoter fragments consist of various promoters that can be chosen based on cell type and desired expression levels. N’ terminus and C’ terminus fragments consist of functional domains that enable various CRISPR mechanisms, including nuclear localization signals (NLSs) for knockout, transactivation domains for CRISPRa, repression domains for CRISPRi, deaminase domains for base editing, and reverse transcriptase domains for prime editing. Cas protein fragments encompass Cas enzymes from different species, including deactivated and nickase versions, as well as Cas variants that have been engineered to have high fidelity or more flexible PAM requirements. Lastly, selection fragments confer antibiotic resistance or visual markers when expressed in eukaryotic cells; these are separated from the Cas protein by the use of 2A sites or internal ribosome entry site (IRES) elements.

**Figure 1.**
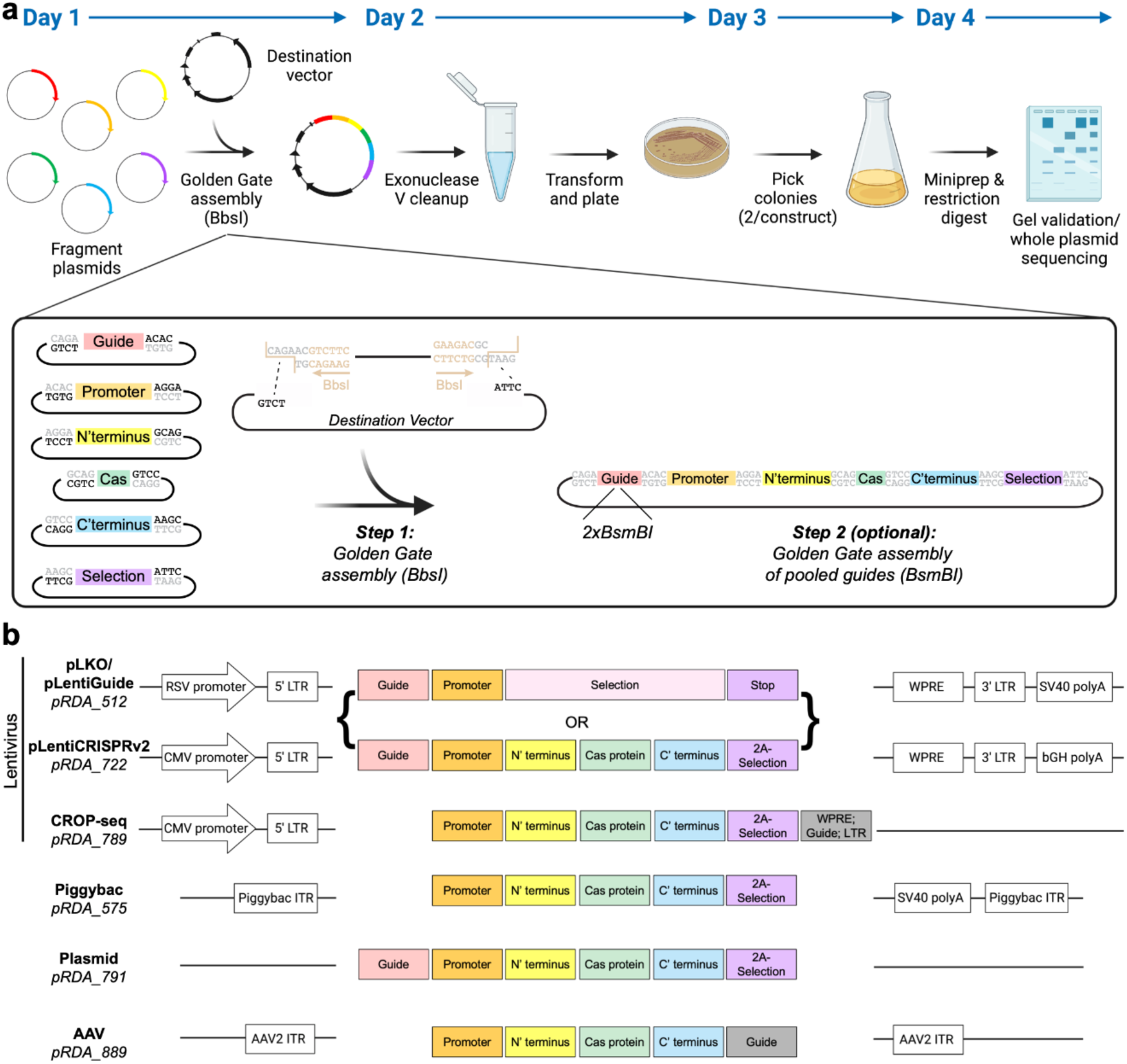
Fragmid overview. **A**) Schematic overview and timeline of the Golden Gate cloning approach. Individual modules and a sample ligation are depicted. **B**) Schematic depicting available destination vectors for various delivery methods, along with compatible inserts. All destination vectors contain an ampicillin resistance cassette and an origin of replication, which are not schematized.

The set of six fragment types is compatible with multiple destination vectors, enabling delivery via lentivirus, PiggyBac-based transposition, and AAV, as well as a minimal plasmid destination vector containing only ampicillin resistance and a replication origin (Figure 1b). Additional destination vectors are readily enabled via repurposing of existing fragments and the use of additional modules. For example, for single-cell readout approaches like CROP-seq^14,15^, a new destination vector and new fragment type were designed to move the guide cassette into the 3’ LTR while maintaining compatibility with existing fragment types. Similarly, due to size constraints for AAV delivery, the 2A-selection fragment is omitted. These modular destination vectors and fragments can be mixed and matched via Golden Gate cloning to create CRISPR vectors that are compatible with numerous enzymes, mechanisms, and delivery modalities.

### Creating constructs

Fragmid utilizes a Type IIS restriction enzyme, BbsI, that cleaves outside of its recognition sequence, allowing for unique, customizable four base pair overhangs, which we refer to as modules (Supplementary Figure 1a). All source fragments have two of these modules, one on each side of the insert, that allow for unidirectional assembly into a destination vector. These four base pair modules are indicated by a number, thus each fragment type and destination vector is defined by two numbers. These modules were chosen using the NEBridge Ligase Fidelity Viewer^16^, resulting in a compatible set of BbsI overhangs predicted to yield 99% correctly-ligated products during GG assembly. All fragments and destination vectors are scrubbed of excess BbsI and BsmBI sites, the latter of which is reserved for subsequent cloning of custom individual guides or guide libraries. Fragments contain the kanamycin resistance cassette, while destination vectors contain the ampicillin resistance cassette, enabling selection for correct assemblies using ampicillin.

For most assembled constructs, the translational start codon appears in the N’ terminal fragment and the stop codon in the 2A-selection fragment, and thus the modules connecting the protein coding fragment types (N’ terminal, Cas, C’ terminal, and 2A-selection) were chosen specifically to code for glycine-serine linkers to minimize disruption of protein properties (Supplementary Figure 1b). Note that in cases where expression of a Cas protein is not desired – for example, to create a vector that expresses only a guide – the N’ terminal, Cas, and C’ terminal fragments can be replaced by a Selection fragment while the 2A-selection fragment can be used to deliver either a second selection marker or a small “filler” comprising little more than a stop codon. Likewise, for a vector that only expresses a Cas protein but no guide, a filler fragment can be used in the Guide cassette position.

To assemble constructs, destination vectors are first digested with BbsI and gel extracted, a step that reduces background colonies, and then mixed with insert fragments in a one-pot GG reaction using BbsI and T4 DNA ligase (Figure 1a). Reaction products are treated with exonuclease to degrade any remaining linear DNA then transformed via heat shock. Extracted plasmid DNA from two colonies per construct is verified via restriction digest and further confirmed via commercial whole-plasmid sequencing. We routinely complete this entire pipeline in four days.

To estimate process fidelity, we collected data from 60 individual assemblies over a five month period (Supplementary Figure 1c). 120 clones (2 per assembly) were first assessed via restriction digest and 112 of these passed visual inspection of the resulting banding pattern (93%). Of assemblies with a correct restriction digest, 1 or 2 clones were sent for nanopore-based whole-plasmid sequencing, for a total of 82 clones, of which 80 showed a perfect match to the anticipated map (98%). One of the two clones that failed sequencing was an AAV vector containing a small deletion in one of the inverted terminal repeats (ITR) hairpins, a common failure mode. The second failed clone appeared to contain a small deletion, which may have been an artifact of the nanopore sequencing, a failure mode that is likely to decrease further in frequency as methods to decrease systematic errors in base calling are developed^17^. Notably, the two clones that failed sequencing QC were from different assemblies, so all 60 attempted assemblies yielded at least one fully correct construct on the first try (100%).

### Establishing positive control guides across CRISPR technologies

The initial description of most advances in CRISPR technology occurs in a small number of easy-to-use systems such as the 293T or K562 cell lines, yet the number of systems to which researchers wish to apply these tools is vast. As a particular CRISPR tool or delivery system does not always perform as hoped for in a new cell model, validated positive controls are an important resource for rapidly assessing experimental feasibility. We thus sought to identify a suite of positive control guides for various CRISPR enzyme and technology combinations that are broadly applicable to different model systems and easy to assay. We chose to target endogenous cell surface markers so that efficiency across a population could be readily measured cell-by-cell using flow cytometry.

Using data from the Cancer Dependency Map (DepMap)^18^, we chose 13 cell surface genes that, upon knockout, are generally non-essential across hundreds of cancer cell lines and vary in RNA expression levels, enabling the assessment of both up- and down-regulating technologies (Supplementary Figure 2a). We tiled guides across the promoter region and gene body to enable assessment of knockout, CRISPRa, and CRISPRi technologies, creating two tiling libraries, one for SpCas9 and one for EnAsCas12a; note that screens conducted with the latter have been recently reported for the optimization of CRISPRa technology^19^.

As an example of the experimental pipeline, we conducted a knockout screen to find active SpCas9 guides targeting CD47. We used an A375 (melanoma) cell line engineered to express SpCas9, then transduced these cells with the tiling library in duplicate (Figure 2a). After selection with puromycin, cells were immunostained and sorted for CD47 expression levels on day 7 post-transduction; a representative example of the gating strategy is provided (Supplementary Figure 2b). After sample processing and sequencing, we calculated the fold change between the log-normalized read counts of guides (LFC) in the low and high expressing populations. Replicate LFCs were modestly correlated, with good enrichment of CD47-targeting guides (Figure 2b).

**Figure 2.**
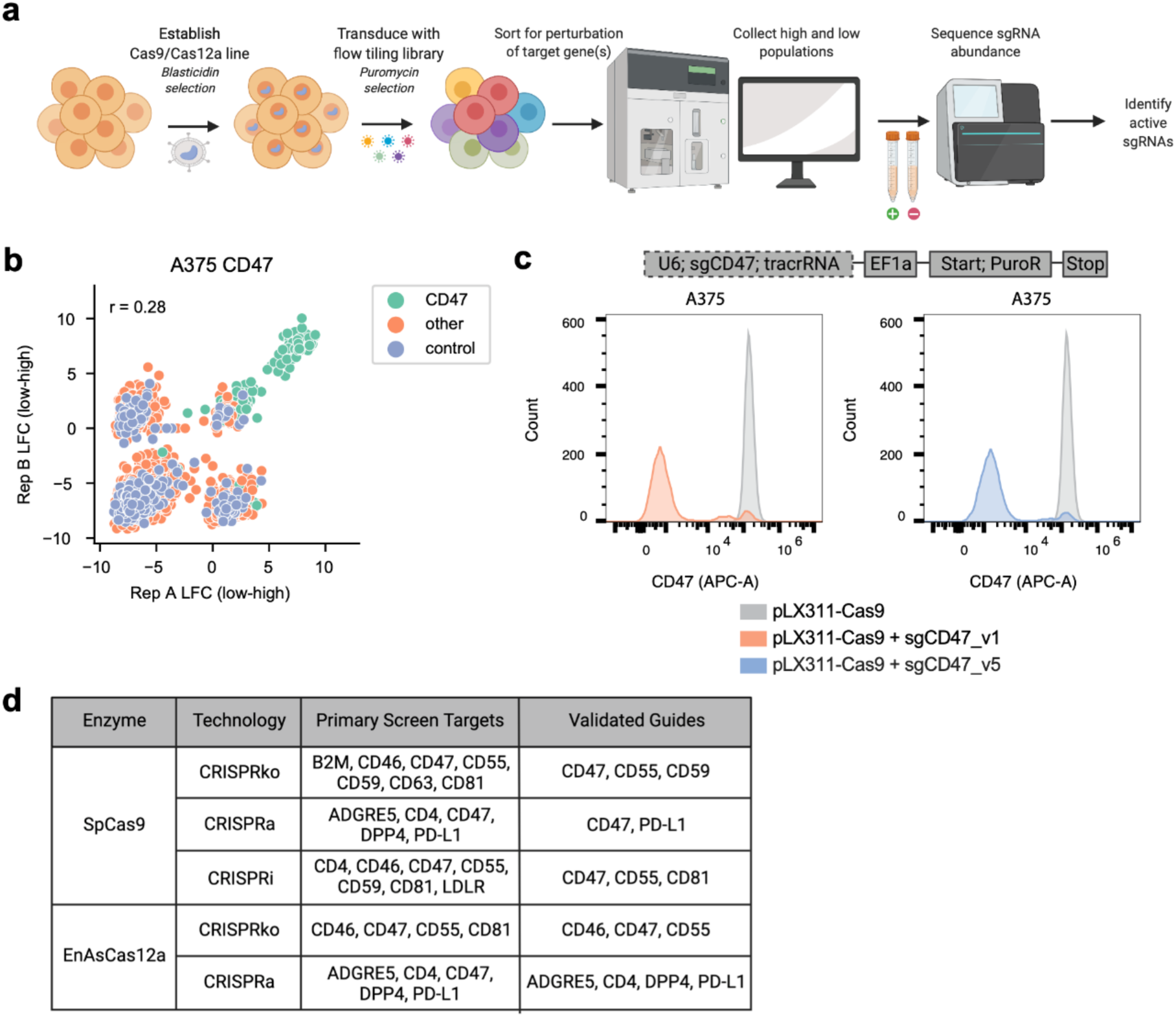
Tiling screens for identification of positive control guides targeting cell surface genes. **A**) Schematic depicting flow cytometry-based tiling screens performed to identify positive control guides. **B**) LFC correlations (Pearson’s r) of the CD47 flow-sorted samples for SpCas9 knockout comparing two replicates. LFCs were calculated by subtracting sorted samples from one another (CD47-negative - CD47-positive sorted population). **C**) Schematic depicting GG-assembled CD47-targeting construct architecture. Dashed lines indicate variable fragment (top). Histograms showing expression levels of CD47 (APC channel) in A375 cells expressing SpCas9 when targeted individually by CD47 guides (bottom). Data from one representative replicate shown. **D**) Table depicting genes targeted in tiling screens and validation experiments for each enzyme-technology combination.

We next sought to validate the top-performing guides from this pooled screen individually. We chose two of the most active CD47-targeting guides from the primary screen and created GG-compatible guide cassette fragments, which were then cloned into constructs containing the EF1a promoter and a puromycin resistance cassette, according to the GG pipeline described above (Figure 2c). These constructs were transduced individually into A375 cells expressing SpCas9, selected on puromycin, and assessed for CD47 expression levels via flow cytometry to confirm knockout activity. Both CD47-targeting guides led to a clear decrease in CD47 expression, and thus were added to the Fragmid toolkit as positive controls for SpCas9 knockout.

The A375 cells expressing the SpCas9 tiling library were then stained and sorted for six additional cell surface targets (B2M, CD46, CD55, CD59, CD63, and CD81) according to the same pipeline (Figure 2d). Guides targeting CD55 and CD59 were chosen for further validation and added to the Fragmid toolkit, while active guides targeting other genes could be easily added to the toolkit in the future, following validation of the primary screen. Similar screens were then performed in A375 and HT29 cells to identify positive control guides for SpCas9 CRISPRa, SpCas9 CRISPRi, EnAsCas12a knockout, and EnAsCas12a CRISPRa (EnAsCas12a CRISPRi yielded poor activity in our hands, although we anticipate testing recent advances^20^). Overall, 28 gene-technology combinations were assessed in this pooled format (Figure 2d). Many top-performing guides were then assessed individually in arrayed format and 15 validated guides were added to Fragmid (Supplementary Figure 2c, Supplementary Data 1).

### Example applications in viral delivery

Making head-to-head comparisons across technologies developed by different research groups, even when plasmids are publicly available, often presents a considerable logistical challenge due to small but potentially important differences in vector design. For example, early genome-scale pooled screening with CRISPR technology used the all-in-one lentiCRISPRv1 vector^21^, which was derived from pLKO, a lentiviral vector widely used for RNAi experiments. The much larger cargo size led to low lentiviral titers and thus several modifications were pursued to generate lentiCRISPRv2^22^. In addition to modification of the cargo, such as the removal of an NLS, a different lentiviral backbone was employed, including differences in the constitutive promoter used upstream of the 5’ LTR and the polyA sequence downstream of the 3’ LTR; lentiCRISPRv1 uses the RSV promoter and SV40 polyA, respectively, as does lentiGuide, while lentiCRISPRv2 utilizes the CMV promoter and bGH polyA. As many progeny vectors for CRISPR technology have subsequently been derived from both lentiGuide and lentiCRISPRv2, we sought to assess the effects of these backbone differences on both lentiviral titer and activity.

To test the performance of lentiGuide and lentiCRISPRv2, we first created GG-compatible destination vectors using the different lentiviral backbone components. We then assembled the same set of fragments into each destination vector to create two all-in-one EnAsCas12a knockout vectors targeting cell surface markers CD47 and CD63 (Figure 3a). A375 and MelJuSo (melanoma) cells were transduced with these lentiviruses in triplicate, and the transduction efficiency was assessed by puromycin selection. In both cell lines, the construct assembled in the lentiCRISPR_v2 destination vector had much higher titer than the construct assembled in the lentiGuide destination vector (Figure 3a). We then assessed the puromycin-selected cells for CD47 and CD63 expression via flow cytometry. Despite the differences in titer, we observed that, once selected with puromycin, the two lentiviral backbones led to similar knockout efficacy (Supplementary Figure 3a).

**Figure 3.**
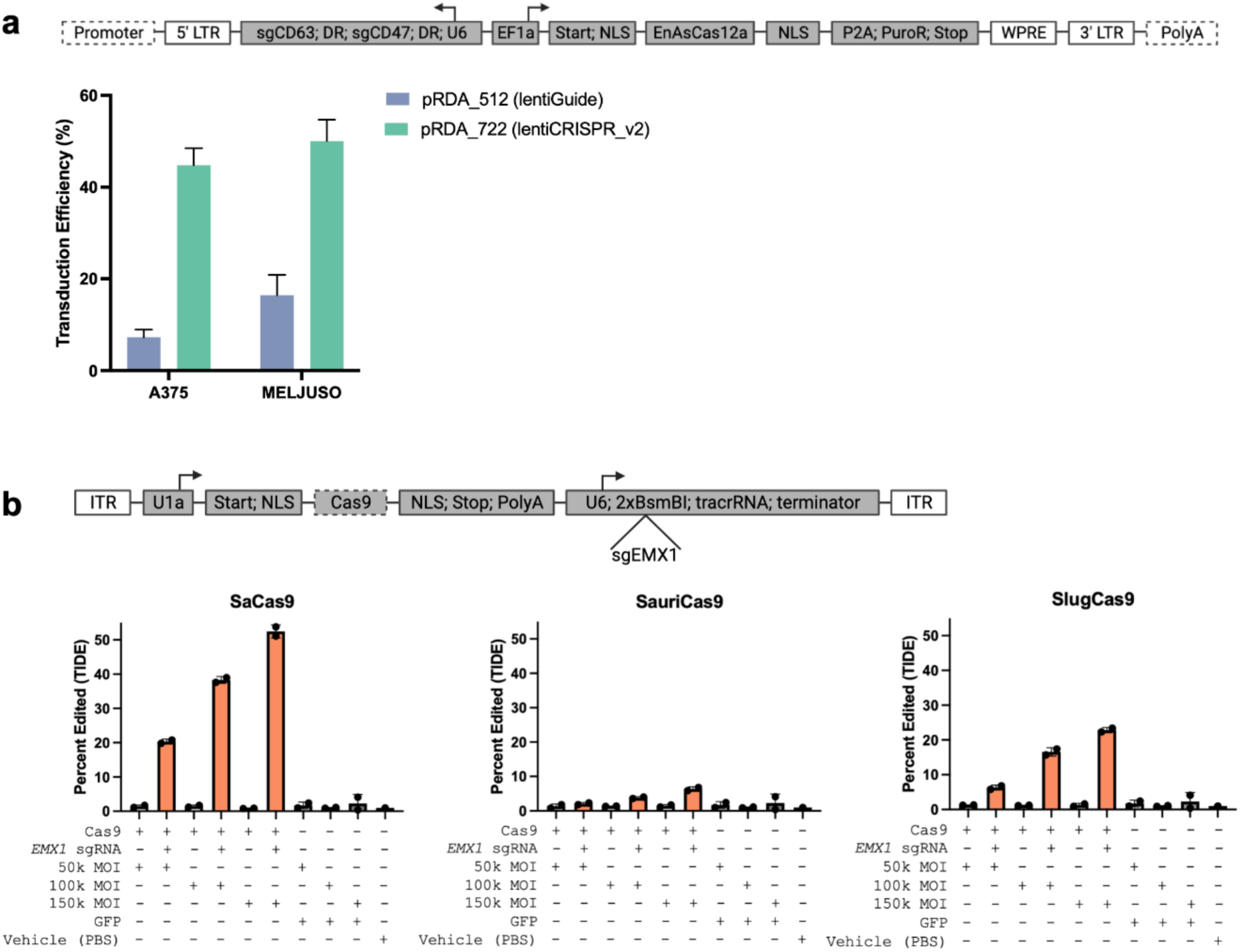
Applications for viral delivery. **A**) Schematic depicting GG assembly of all-in-one EnAsCas12a constructs into the lentiGuide and lentiCRISPR_v2 destination vectors. Dashed lines indicate variable elements in destination vector (top). Barplots illustrating percent of A375 and MelJuSo cells successfully transduced for each construct (bottom). **B**) Schematic depicting GG assembly of all-in-one Cas9 constructs into the AAV destination vector. Dashed lines indicate variable fragment (top). Barplots illustrating editing efficiency, quantified by TIDE, in HEK293T cells transduced with SaCas9, SauriCas9, and SlugCas9 constructs at increasing MOIs, with and without the *EMX1* guide (bottom).

This platform is particularly helpful for the head-to-head comparison of different fragments, such as Cas proteins. Adeno-associated virus (AAV) has become an increasingly popular approach for the *in vivo* delivery of gene therapies and CRISPR-Cas9 editing systems^23–26^. However, AAV has a packaging size limit of <5 kb^27,28^, which presents challenges for encoding CRISPR components. The well-validated *Streptococcus pyogenes* Cas9 (SpCas9) has an open reading frame of 4104 base pairs (bp) which leaves little room to encode an sgRNA within the same AAV vector. Other Cas9 orthologs, including *Staphylococcus aureus* Cas9 (SaCas9) and *Neisseria meningitidis* 2 Cas9 (Nme2Cas9), have shorter open reading frames (3159 bp and 3246 bp, respectively) that can be encoded into all-in-one AAV constructs^29,30^. All-in-one AAV delivery is desirable because it can reduce manufacturing burden and may improve safety by minimizing the viral dose, as high AAV doses are known to cause toxicity^31,32^.

The shorter open reading frame of SaCas9 compared to Nme2Cas9 allows for greater flexibility when used in all-in-one AAVs, and short open reading frames are particularly important when designing all-in-one constructs with multiple sgRNAs^28^. Despite this advantage, SaCas9 has a fairly restrictive PAM requirement of 5’ NNGRRT 3’ (where N is any nucleotide and R is A or G)^29^, while Nme2Cas9 has a less restrictive dinucleotide PAM (5’ NNNNCC 3’), allowing for higher targeting density. Because of these drawbacks, both enzymes are important in the all-in-one AAV toolkit, and selected for experiments depending on use case. An ideal Cas9 for all-in-one AAV delivery would have both features - a small open reading frame and a short PAM sequence. *Staphylococcus auricularis* Cas9 (SauriCas9) and *Staphylococcus lugdunensis* Cas9 (SlugCas9) were recently discovered through their homology to SaCas9. These two enzymes have similar open reading frame sizes to SaCas9 (SauriCas9: 3180 bp; SlugCas9: 3162 bp), but have a more frequently occurring dinucleotide PAM (5’ NNGG 3’)^33,34^, which could make them useful additions to the growing Cas9 all-in-one AAV toolkit.

The ability to quickly assemble and test all-in-one AAVs is critical due to the fact that the orientation of different genes within an AAV genome can affect the quality of the vector genome and thus editing efficiency^35–37^. As SaCas9, SauriCas9, and SlugCas9 utilize the same tracrRNA, we sought to perform a head-to-head comparison of these three enzymes using the GG assembly platform. We designed an AAV destination vector that is compatible with the GG pipeline, and uses most of the same fragment types as before (Figure 1b). Three all-in-one constructs containing SaCas9, SauriCas9, and SlugCas9 were assembled into the AAV destination vector via the GG pipeline (Figure 3b). A guide sequence targeting the human *EMX1* locus that was previously shown to be edited by all three Cas9s of interest^33,34^ was then cloned into these three constructs via BsmBI in a subsequent step.

All constructs, with and without the *EMX1* guide cloned into the sgRNA, were then used to produce AAV6 particles. We also included a pAAV-GFP plasmid, which encodes GFP only and was not assembled via the GG pipeline, to serve as both a control for the assembly pipeline and a negative control for downstream editing experiments. The resulting AAV6 viruses were purified and then quantified using droplet digital PCR (ddPCR) with primers specific to the ITR sequence of AAV to determine the concentration of viral preparations. Viral titers were similar for all viruses produced, indicating that the Golden Gate assembly process did not negatively affect the generation of AAV particles (Supplementary Figure 3b). We then transduced HEK293T cells in duplicate with either a vehicle control (PBS) or with increasing multiplicities of infection (MOIs) for the purified viruses. 72 hours after viral transduction, genomic DNA was collected from all samples, the *EMX1* locus was PCR amplified and Sanger sequenced, and editing was quantified using Tracking of Indels by DEcomposition (TIDE)^38^. As expected, all three of the GG-assembled all-in-one viruses containing the *EMX1* sgRNA edited the HEK293T cells in an MOI-dependent manner, indicating their functionality (Figure 3b). SaCas9 yielded the highest editing efficiency at *EMX1* when transduced at the highest MOI (52% edited) followed by SlugCas9 and SauriCas9 (23% and 6% edited, respectively). While neither of the recently-identified enzymes performed better than SaCas9 at this locus, they are still useful additions to the all-in-one AAV toolkit, as their different PAM requirements may be preferable for some use cases.

### Higher throughput assessment of vector elements

The aforementioned examples of head-to-head comparisons required individual assembly of various constructs, but the modular nature of this approach also lends itself to pooled screens. We attempted to take a hybrid approach in which constructs were GG assembled in arrayed format, including plate- and well-specific barcodes, but then all subsequent steps were performed in a pool (Supplementary Figure 4a). This hybrid approach provides many of the benefits of a pooled screen, including removing the need for liquid handling robots to manipulate mammalian cells, as well as the internally-controlled conditions inherent to a pooled screen. We tested this strategy by comparing 3 different down-regulation technologies in which the Cas protein of interest was driven by four different promoters. We compared SpCas9-based knockout and interference as well as EnAsCas12a-based knockout, assaying down-regulation of a cell surface marker by flow cytometry and depletion of essential genes via viability screens.

The three essential genes we chose to target were *DBR1*, *RPL3*, and *PLK1*, nominated both by their performance in SpCas9 knockout screens in the DepMap and via social media as fast-killing target genes, while *CD47* was chosen due to its ubiquitous expression across commonly used cell lines and its non-essentiality in cell culture. We assembled the relevant fragments in a modified lentiviral destination vector that allowed the addition of barcodes downstream of the stop codon in the 2A-selection fragment, so that the assembled constructs could be uniquely identified via short Illumina reads. Each combination of promoter (4), Cas activity type (3), and corresponding guide (6: 3 essential guides, 1 *CD47* guide, and 2 intergenic control guides) were assembled in 4 replicate wells, each with its own barcode, for a total of 288 constructs. These plasmid DNAs (pDNAs) were pooled post-GG assembly and electroporated into *E. coli.* Subsequent sequencing of the amplified pDNA via the barcode showed a good distribution of clones (AUC = 0.64, Supplementary Figure 4b) which falls in the typical range for guide libraries (AUC values range from 0.6 - 0.7). There is much more variation in construct size in this pool than in a standard pooled guide library, as the longest and shortest constructs differ by 1.3 kb, so this AUC value indicates the high fidelity of this assembly approach.

The library was screened in duplicate in MelJuSo cells. We first conducted a screen in which we sorted the cells for CD47 levels, collecting the top 50% and bottom 25% of the population. We also conducted a viability screen in which an early time point was taken at day 4 post-transduction and a late time point at day 14. Importantly, for the viability screen, we used lentivirus that was treated with benzonase to avoid plasmid contamination of the early time point sample that would otherwise misrepresent the true abundance of each construct in the lentiviral pool due to unequal packaging efficiency of constructs of different length^39^. Pearson correlation of LFC values across replicates for the CD47 sort screen was 0.67 and for the viability screen 0.88 (Supplementary Figure 4c).

In the CD47 screen, we observed excellent specificity for all three technologies, with guides targeting *CD47* clearly enriching in the low expression bin relative to intergenic controls (Figure 4a). Likewise, in the viability arm, we saw depletion of the constructs targeting the essential genes with all three technologies relative to intergenic controls. Importantly, we observed little variation across the 4 barcodes assigned to each construct. This observation mitigates a substantial concern we had with this approach, namely that shuffling of lentiviral elements during reverse transcription would lead to substantial decoupling^40^ between the barcode and the targeting guide, which are separated by several thousand base pairs. While quantitating the degree of uncoupling is technically challenging in this experimental design, the demonstrated performance of these screens suggests that the magnitude of this confounding phenomenon does not obscure all biological signal. We observed that different technologies performed similarly regardless of the Pol II promoter used to express the Cas protein, with EF1a performing slightly better across most technologies and targets (Figure 4b). Although it would be inappropriate to conclude that one technology is fundamentally better based on the results of this experiment, as each is assessed with only four guides and in one cell line, such pools comprising diverse technologies represent a useful resource for rapidly nominating a particular approach for a cell model of interest.

**Figure 4.**
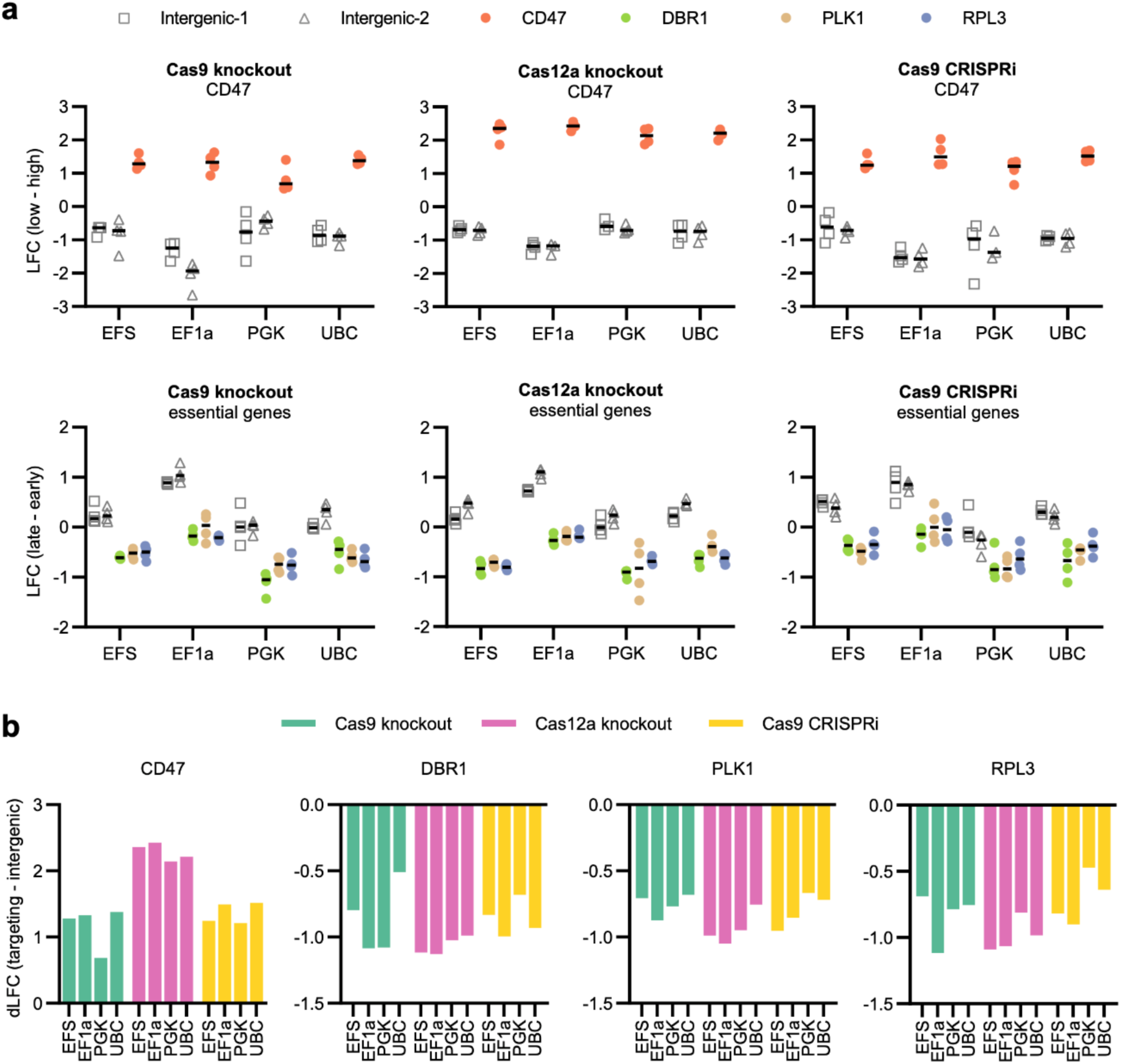
Pooled Golden Gate assembly and assessment. **A**) LFC values (CD47-low - CD47-high) binned by technology and promoter (top). LFC values (late-early time points) for the viability screen in MelJuSo cells, binned by technology and promoter (bottom). Each point represents a unique barcode; black line indicates the median. **B**) dLFC values (targeting - intergenic) for the CD47-sort screen (left) and viability screen (right), binned by targeted gene and promoter.

### Assembly of plasmids for *Drosophila* cell-based and in vivo expression

Because of its rapid life cycle, and the ease of generating transgenics, *Drosophila* is well suited as an in vivo system for rapid assessment of new Cas variants and fusion proteins. The recent development of CRISPR pooled screening in *Drosophila* cells has further expanded the types of genome-wide screens that are now possible^41–44^. To extend the Fragmid toolkit described above to *Drosophila*, several destination vectors, guide cassettes, and promoter fragments were produced that are compatible with the modular assembly and suitable for generating transgenic flies or stable cell lines. Test plasmids were assembled for each of the three *Drosophila* destination vectors, each expressing a Cas protein and fluorescent marker. The assembled fragments were correctly expressed under the control of the *Drosophila* promoters, and the destination vectors produced the expected germline transformation or integration in cells (Supplementary Figure 5).

### Fragmid overview

The Fragmid ecosystem is enabled via a web portal: http://broad.io/fragmid. Users first select from available organisms (currently mammalian or *Drosophila*), delivery types (i.e. destination vectors), CRISPR enzymes (i.e. Cas proteins), and CRISPR mechanisms (i.e. genome engineering applications). Based on these inputs, compatible fragments are filtered and displayed, which users may then browse and select from. Fragmid then performs an *in silico* assembly of the desired construct and delivers a .gb file of the assembled map to the user. We emphasize that the Fragmid system enables mixing and matching of many elements, and an incomplete understanding of the functionality of different components may result in nonsensical vectors, so careful design consideration by the user remains necessary when using this toolkit.

Available guide cassettes (flanked by BbsI modules 0 and 1) are determined by target type, CRISPR enzyme, and CRISPR mechanism. As described above, positive control fragments include validated control guides that are specific to each Cas-mechanism combination, while cloning fragments include BsmBI sites for subsequent introduction of custom individual guides or pooled libraries. The most common RNA polymerase III cassette for guide expression is the human U6 promoter^45,46^, while the H1 promoter is also occasionally used and is included here for dual guide expression^47^. There are also numerous tracrRNA variants for SpCas9, which include modifications to increase RNA expression levels^48,49^, enable the use of unique molecular identifiers (UMIs)^50^, and recruit proteins via PP7 and MS2 aptamers^51–53^. For expressing Cas12a guides, note that vectors that also include the Cas12a protein (all-in-one vectors) require reversing the Pol III promoter if using lentivirus, otherwise the primary lentiviral transcript will be cut by Cas12a in the 293T packaging cells. Cas12a guide fragments are therefore available in the reverse direction, defined as having the Pol III promoter point towards the 5’LTR instead of the 3’LTR. Finally, in cases in which guide expression is not desired, such as the creation of a Cas-only vector, a small filler sequence can be used in this position.

Cas protein options (modules 3 and 4) are determined by both the CRISPR enzyme and CRISPR mechanism chosen. For knockout, nuclease versions of SpCas9, SaCas9, SlugCas9, SauriCas9, AsCas12a, and EnAsCas12a are available, as well as PAM-flexible engineered SpCas9 variants SpRY, SpG, and NG^54,55^. For CRISPRa and CRISPRi, nuclease-deactivated versions of many of these enzymes are available. Similarly, D10A and H840A nickase versions of SpCas9 are available for base editing and prime editing applications, respectively. RNA-targeting Cas proteins are included as well: RfxCas13d^56^, DjCas13d^57^, and Cas7-11^58^. Finally, note that the same module used for Cas proteins can instead be repurposed to generate fusion proteins, with either the PP7 core protein (PCP) or MS2 core protein (MCP) for use with modified tracrRNAs as discussed above, or with the ALFA nanobody for use with the ALFA tag^19^.

The functional domains available in the N’ terminus (modules 2 and 3) and C’ terminus (modules 4 and 5) fragments are displayed based on the selected CRISPR mechanism. For knockout, one or more NLSs are recommended at each termini, of which there are several options, both monopartite and bipartite. For CRISPRa, several common transactivation domains, including VP64, p65, HSF1, and p300 are available, while for CRISPRi, repression domains include the KRAB domains from ZNF10 (also known as KOX1) and ZIM3, as well as MeCP2. Deaminase and reverse transcriptase domains are available for base editing and prime editing, respectively.

Pol II promoters (modules 1 and 2) are agnostic to the enzyme and mechanism chosen, and should be selected based on vector size limitations, cell type, and desired readout approach, among other experimental design constraints. Fragmid includes several widely-used constitutive Pol II promoters, including EF1a (human elongation factor-1 alpha), PGK (human phosphoglycerate kinase), UbC (human ubiquitin C), and SFFV (spleen focus-forming virus). For delivery mechanisms where cargo size is limited, such as AAV, smaller promoters such as U1a or EFS (human core/short EF1a) may be advantageous, as these are ∼1 kb shorter than some of the above promoters.

Positive selection markers (modules 5 and 6) are also independent of CRISPR enzyme and mechanism, and should be chosen based on desired experimental readout. Here we provide components from two main categories of positive selection markers, antibiotic resistance genes and fluorescent proteins. We provide selection fragments conferring resistance to puromycin, blasticidin, hygromycin, and G418/neomycin, as well as numerous fluorescent proteins such as EGFP, mCherry, TagRFP, TagRFP657, mTagBFP, and VexGFP. Most selection cassettes are expressed via the use of 2A sites^59,60^ but there are also fragments that contain IRES elements^61,62^ that allow for independent translation of downstream selection markers, and thus are preceded by a stop codon. In some circumstances, such as the creation of a guide-only vector, a central Cas protein is not desired and in these cases one can omit inclusion of the 2-3, 3-4, and 4-5 modules and instead use a selection cassette with a 2-5 module. A second selection marker could then be added with the 5-6 module, or a small filler sequence used instead.

Importantly, Fragmid incorporates versioning control to help distinguish between components that may share a common name but have distinct nucleotide sequences. For example, there are several tracrRNA variants for SpCas9 provided, which are displayed as trRNA_v1, trRNA_v4, etc. Note that the incremented version is not meant to indicate that prior versions are now deprecated, but rather simply reflects the order in which the specific component was added to the Fragmid system; versions that do not appear on the web portal often represent components that were tested internally but, for any number of scientific or organizational reasons, were not selected for general usage. For protein-coding sequences, two version numbers are applied, the first for the amino acid sequence and the second for the nucleotide sequence. For example, KRAB_v1.1 and KRAB_v2.1 represent the KRAB domains from ZNF10 and ZIM3, respectively, which have different amino sequences. In contrast, VP64_v1.1 and VP64_v1.2 are comprised of the same amino acid sequence but differ in their underlying nucleotide sequence, in order to mitigate the potential for recombination if included on the same vector, such as including a copy at both the N’ and C’ termini of a Cas protein.

## DISCUSSION

Here we develop Fragmid, a modular toolkit with standardized interchangeable parts, to accelerate the use of CRISPR technologies. This resource comprises a repertoire of destination and fragment vectors that span many delivery modalities, enzymes, and mechanisms. Further, we develop a suite of positive control guides to enable facile benchmarking when testing CRISPR technology in a new experimental setting or when developing new approaches. We then utilize the toolkit to rapidly create and test vectors for a variety of applications, including improving lentiviral titer and testing new enzymes for all-in-one AAV delivery. Finally, we create a public online portal to make this toolkit approachable.

Fragmid also enabled the assessment of multiple CRISPR technologies simultaneously in a pooled setting. This has the potential to rapidly accelerate the credentialing of new model systems for subsequent large-scale screens. While here we tested three down-regulation approaches – knockout with SpCas9 and EnAsCas12a, and CRISPRi with SpCas9 – analogous pools could assess and optimize other technologies, for example, to understand what combination of activators works best in a particular cell model or at a particular gene, or which RNA-targeting enzymes, if any, provide sufficient on-target activity while avoiding non-specific collateral activity when stably expressed. Further, we envision that alternative barcoding strategies and long-read sequencing could enable such resources to be assembled using an entirely pooled approach.

Especially in a non-profit environment, every large-scale resource runs the risk of near-instantaneous obsolescence following initial publication. The first challenge is uptake by potential users, that is, research groups looking to use CRISPR technology to answer biological questions of interest. We hope that the ability to conveniently test numerous approaches in parallel will prove valuable, rather than the traditional approach of acquiring one or two vectors from a nearby lab and hoping that it works well-enough (an often hazily-defined performance metric) in a new experimental setting. Standardized positive controls should be beneficial, enabling controlled, head-to-head optimization. A second challenge is that CRISPR technology continues to evolve rapidly. Our hope is that labs developing new approaches will see the value in using this system for testing new Cas proteins and domains to accelerate their own explorations (Supplementary Note 1). We anticipate that deposition of new fragments with Addgene and incorporation into the Fragmid portal will allow for faster dissemination of new technologies, maximizing their impact beyond the initial publication.

## ACKNOWLEDGEMENTS

We thank all members of the Genetic Perturbation Platform (GPP); Desiree Hernandez, Monica Roberson, Eliezer Josue Ibarra, Pema Tenzing, Tashi Lokyitsang, and Xiaoping Yang for producing guide libraries and lentivirus; the Broad Institute Genomics Platform Walk-up Sequencing group for Illumina sequencing; and Ganna Reint for helpful discussions. This work was supported in part by NCI U01CA250565 to JGD and the Functional Genomics Consortium, a partnership between the GPP at the Broad Institute, Merck, Janssen, Vir, Abbvie, and Bristol Meyers Squibb. Development and validation of *Drosophila* components at the Drosophila Research & Screening Center-Biomedical Technology Research Resource (DRSC-BTRR) was supported by NIGMS P41 GM132087 to Stephanie E. Mohr and Norbert Perrimon, and the DRSC-BTRR thanks Karim Rahimi and Yousuf Hashmi for input on the project.

## AUTHOR CONTRIBUTIONS

Conceptualization: AVM, JGD

Investigation: AVM, YVL, ALG, ZMS, CK, NWM, RJS, BEV, AG, JAB, JDZ, RV

Analysis: AVM, YVL, ZMS

Data Curation: AVM, ZMS, YVL

Software: BW, MAG, TMG

Visualization: AVM, RJS, CK, ZMS

Supervision: EJS, JGD

Writing: AVM, CK, JDZ, JGD

## METHODS

### Vectors

All destination vectors and fragments included in the first release of Fragmid are available on the web portal (http://broad.io/fragmid) and will be deposited with Addgene.

pAAV-CAG-GFP was a gift from Karel Svoboda (Addgene plasmid #28014).

pAAV2/6 was generously provided by Dr. Guangping Gao and Dr. Jun Xie of the UMass Chan Medical School Viral Vector Core.

pAdDeltaF6 was a gift from James M. Wilson (Addgene plasmid #112867).

### Cell lines and culture

A375, HCT116, HT29, and MelJuSo cells were obtained from the Cancer Cell Line Encyclopedia at the Broad Institute. HEK293T cells were obtained from ATCC (CRL-3216).

All cells regularly tested negative for mycoplasma contamination and were maintained in the absence of antibiotics except during screens, flow cytometry-based experiments, and lentivirus production, during which media was supplemented with 1% penicillin-streptomycin. Cells were passaged every 2-4 days to maintain exponential growth and were kept in a humidity-controlled 37°C incubator with 5.0% CO2. Media conditions and doses of polybrene, puromycin, and blasticidin were as follows, unless otherwise noted:

A375: RPMI + 10% fetal bovine serum (FBS); 1 μg/mL; 1 μg/mL; 5 μg/mL

HCT116: McCoy’s 5A + 10% FBS; 4 μg/mL; 2 μg/mL; 16 μg/mL

HT29: DMEM + 10% FBS; 1 μg/mL; 2 μg/mL; 8 μg/mL

MelJuSo: RPMI + 10% FBS; 4 μg/mL; 1 μg/mL; 4 μg/mL

HEK293T: DMEM + 10% heat-inactivated FBS; N/A; N/A; N/A

### SpCas9 cell surface marker tiling library design

Guide sequences for the tiling library were designed using sequence annotations from Ensembl (GRCh38). CRISPick was used to select every possible guide (using an NNNN PAM) against the longest annotated transcript for 17 genes: CD47, CD63, B2M, CD274, CD46, CD55, CD81, CSTB, CD4, CD26, CD97, CD59, BSG, LDLR, LRRC8A, PIGA, and TFRC. We included guides targeting the coding sequence, all guides for which the start was up to 30 nucleotides into the intron and UTRs, and all guides targeting the window 0-300 bp upstream of the annotated transcription start site (TSS). The library was filtered to exclude any guides with BsmBI recognition sites or TTTT sequences, and guides were annotated to denote the CRISPR technologies with which they were compatible (knockout, CRISPRa, CRISPRi and/or base editing). Guides with >3 or >5 perfect matches in the genome for knockout/base editing or CRISPRa/CRISPRi technologies, respectively, were also filtered out. 700 positive and negative control guides were added into the library, including 500 guides targeting intergenic regions, 100 non-targeting guides, and 100 guides targeting essential splice sites, for a total library size of n = 6,605. Guides targeting genes used for viability and coselection screening purposes as well as base editing guides were not assessed in this study.

### EnAsCas12a cell surface marker tiling library design

Guide sequences for the tiling library were designed using sequence annotations from Ensembl (GRCh38). CRISPick was used to select every possible guide (using an NNNN PAM) against the longest annotated transcript for 17 genes: CD47, CD63, B2M, CD274, CD46, CD55, CD81, CSTB, CD4, CD26, CD97, CD59, BSG, LDLR, LRRC8A, PIGA, and TFRC. We included guides targeting the coding sequence, all guides for which the start was up to 30 nucleotides into the intron and UTRs, and all guides targeting the window 0-300 bp upstream of the annotated TSS. The library was filtered to exclude any guides with BsmBI recognition sites or TTTT sequences, and guides were annotated to denote the CRISPR technologies with which they were compatible (knockout, CRISPRa, CRISPRi and/or base editing). Guides with >3 or >5 perfect matches in the genome for knockout/base editing or CRISPRa/CRISPRi technologies, respectively, were also filtered out. Subsequently, a random 50% subsampling of the knockout/base editing guides was removed from the library to decrease library size. 700 positive and negative control guides were added into the library, including 500 guides targeting intergenic regions, 100 non-targeting guides, and 100 guides targeting essential splice sites, for a total library size of n = 8,421. Guides targeting genes used for viability and coselection screening purposes as well as base editing guides were not assessed in this study.

### Modular vector destination vector pre-digestion

Destination vectors were synthesized and sequence-verified by Genscript. Destination vectors were pre-digested using BbsI (New England Biolabs) at 37°C for two hours. Linearized destination vectors were gel purified using 0.7% agarose gels and extracted with the Monarch DNA Gel Extraction Kit (New England Biolabs), before further purification by isopropanol precipitation.

### Modular vector Golden Gate assembly

Fragments were synthesized, cloned into the pUC57-Kan backbone, and sequence-verified by Genscript. Fragments were diluted to 10 nM and cloned into a pre-digested destination vector in 30 uL Golden Gate cloning reactions. Each reaction contained 3 uL BbsI (New England Biolabs), 1.25 uL T4 ligase (New England Biolabs), 3 uL of 10x T4 ligase buffer (New England Biolabs), 75 ng destination vector, and a 1:1 molar ratio of fragments:destination vector. Reactions were carried out under the following thermocycler conditions: (1) 37°C for 5 minutes; (2) 16°C for 5 minutes; (3) go to (1), x 100; (4) 37°C for 30 minutes; (5) 65°C for 20 minutes. The ligation product was treated with Exonuclease V (New England Biolabs) at 37°C for 30 minutes before enzyme inactivation with the addition of EDTA to 11 mM. Per reaction, 10 uL of product was transformed into Stbl3 chemically competent *E. coli* (Invitrogen) via heat shock, and grown at 37°C for 16 hours on agar with 100 μg/mL carbenicillin. Colonies were picked and grown at 37°C for 16 hours in 5 mL LB with 100 μg/mL carbenicillin. Plasmid DNA (pDNA) was prepared (QIAprep Spin Miniprep Kit, Qiagen). Purified plasmids were sequence confirmed by restriction enzyme digests and whole plasmid sequencing.

### Whole-plasmid sequencing QC

Assembled plasmids were submitted to Plasmidsaurus or Primordium Labs for nanopore-based whole plasmid sequencing. Samples sent to Plasmidsaurus were diluted to 30 ng/uL and sent as 10 uL aliquots. Samples were sent to Primordium as undiluted aliquots of at least 4 uL at concentrations >200 ng/uL. Samples with up to three mismatches from the expected sequence were considered as passing QC, with the exception of any mismatches in BbsI overhang regions, which were considered as failing QC. Additionally, small insertions or deletions in homopolymer tracts were assumed to be artifacts of nanopore sequencing and considered as passing.

### Golden Gate assembly for pooled screens

Fragments were cloned into pre-digested destination vector via Golden Gate cloning in 96-well plates, using a 2:1 molar ratio of fragments:destination vector. A 96-well plate of 7-10 nt barcodes, flanked by 6-9 module BbsI sites, was used to barcode each well of the reaction uniquely. To further enable multiplexing, a unique 7 nt plate barcode was designed for each plate, flanked by 9-10 modules, allowing all three technologies (SpCas9 knockout, SpCas9 CRISPRi, and EnAsCas12a knockout) to be assembled in separate plates then pooled together. The ligation products were treated with Exonuclease V (New England Biolabs) in each 96-well plate, then 5 uL from each well was pooled. The pooled product was then isopropanol precipitated and electroporated into Stbl4 electrocompetent cells (Invitrogen) and grown at 30°C for 16 hours on agar with 100 μg/mL carbenicillin. Colonies were scraped and plasmid DNA (pDNA) was prepared (HiSpeed Plasmid Maxi, Qiagen). To confirm library representation and distribution, the pDNA was sequenced by Illumina HiSeq2500 High Output.

### Library production

Oligonucleotide pools were synthesized by CustomArray. BsmBI recognition sites were appended to each sgRNA sequence along with the appropriate overhang sequences (bold italic) for cloning into the sgRNA expression plasmids, as well as primer sites to allow differential amplification of subsets from the same synthesis pool. The final oligonucleotide sequence for Cas9 systems was thus: 5′-[Forward Primer]CGTCTCA***CACCG***[sgRNA, 20 nt]***GTTT***CGAGACG[Reverse Primer], and for Cas12a systems: 5′-[Forward Primer]CGTCTCA***AGAT***[sgRNA, 23 nt]***GAAT***CGAGACG[Reverse Primer].

Primers were used to amplify individual subpools using 25 μL 2x NEBnext PCR master mix (New England Biolabs), 2 μL of oligonucleotide pool (∼40 ng), 5 μL of primer mix at a final concentration of 0.5 μM, and 18 μL water. PCR cycling conditions: (1) 98°C for 30 seconds; (2) 53°C for 30 seconds; (3) 72°C for 30 seconds; (4) go to (1), x 24.

The resulting amplicons were PCR-purified (Qiagen) and cloned into the library vector via Golden Gate cloning with Esp3I (Fisher Scientific) and T7 ligase (Epizyme); the library vector was pre-digested with BsmBI (New England Biolabs). The ligation product was isopropanol precipitated and electroporated into Stbl4 electrocompetent cells (Invitrogen) and grown at 30°C for 16 hours on agar with 100 μg/mL carbenicillin. Colonies were scraped and plasmid DNA (pDNA) was prepared (HiSpeed Plasmid Maxi, Qiagen). To confirm library representation and distribution, the pDNA was sequenced.

### Lentivirus production

For small-scale virus production, the following procedure was used: 24 hours before transfection, HEK293T cells were seeded in 6-well dishes at a density of 1.5 × 10^6^ cells per well in 2 mL of DMEM + 10% heat-inactivated FBS. Transfection was performed using TransIT-LT1 (Mirus) transfection reagent according to the manufacturer’s protocol. Briefly, one solution of Opti-MEM (Corning, 66.75 μL) and LT1 (8.25 μL) was combined with a DNA mixture of the packaging plasmid pCMV_VSVG (Addgene 8454, 125 ng), psPAX2 (Addgene 12260, 1250 ng)^63^, and the transfer vector (e.g., pLentiGuide, 1250 ng). The solutions were incubated at room temperature for 20–30 min, during which time media was changed on the HEK293T cells. After this incubation, the transfection mixture was added dropwise to the surface of the HEK293T cells, and the plates were centrifuged at 1000 x g for 30 minutes at room temperature. Following centrifugation, plates were transferred to a 37°C incubator for 6–8 hours, after which the media was removed and replaced with DMEM + 10% FBS media supplemented with 1% BSA. Virus was harvested 36 hours after this media change.

A larger-scale procedure was used for pooled library production. 24 hours before transfection, 18 × 10^6^ HEK293T cells were seeded in a 175 cm^2^ tissue culture flask and the transfection was performed the same as for small-scale production using 6 mL of Opti-MEM, 305 μL of LT1, and a DNA mixture of pCMV_VSVG (5 μg), psPAX2 (50 μg), and 40 μg of the transfer vector. Flasks were transferred to a 37°C incubator for 6–8 hours; after this, the media was aspirated and replaced with BSA-supplemented media. Virus was harvested 36 hours after this media change.

### Determination of antibiotic dose

In order to determine an appropriate antibiotic dose for each cell line, cells were transduced with the pRosetta or pRosetta_v2 lentivirus such that approximately 30% of cells were transduced and therefore EGFP+. At least 1 day post-transduction, cells were seeded into 6-well dishes at a range of antibiotic doses (e.g., from 0 μg/mL to 8 μg/mL of puromycin). The rate of antibiotic selection at each dose was then monitored by performing flow cytometry for EGFP+ cells. For each cell line, the antibiotic dose was chosen to be the lowest dose that led to at least 95% EGFP+ cells after antibiotic treatment for 7 days (for puromycin) or 14 days (for blasticidin).

### Determination of lentiviral titer

To determine lentiviral titer for transductions, cell lines were transduced in 12-well plates with a range of virus volumes (e.g., 0, 150, 300, 500, and 800 μL virus) with 3 × 10^6^ cells per well in the presence of polybrene. The plates were centrifuged at 640 x g for 2 hours and were then transferred to a 37°C incubator for 4–6 hours. Each well was then trypsinized, and an equal number of cells seeded into each of two wells of a 6-well dish. Two days post-transduction, puromycin was added to one well out of the pair. After 5 days, both wells were counted for viability. A viral dose resulting in 30–50% transduction efficiency, corresponding to an MOI of ∼0.35–0.70, was used for subsequent library screening.

### Pooled screens

For pooled screens, cells were transduced in two biological replicates with the lentiviral library. To remove residual pDNA contamination, lentivirus was treated with 2000 U/mL Benzonase (Millipore Sigma, Product No. E1014) at 37°C for 30 minutes immediately prior to transduction, in a buffer consisting of 50 mM Tris HCl (pH 8.0, Millipore Sigma, Product No. T2694-100ML), 1 mM MgCl2 (Millipore Sigma, Product No. M8787) and 100 μg/mL BSA (Millipore Sigma, Product No. A3294-10G)^39^. Transductions were performed at a low multiplicity of infection (MOI ∼0.3), using enough cells to achieve a representation of at least 1000 transduced cells per guide, assuming a 30% transduction efficiency. Cells were plated in polybrene-containing media with 3 x 10^6^ cells per well in a 12-well plate. Plates were centrifuged at 821 x g for 2 hours, after which 2 mL of media was added to each well. Plates were then transferred to an incubator for 4-6 hours, after which virus-containing media was removed and cells were pooled into flasks. Puromycin was added 2 days post-transduction and maintained for 5 days to ensure complete removal of non-transduced cells. Upon puromycin removal, cells were passaged every 2-3 days for an additional 1 week to allow guides to enrich or deplete; cell counts were taken at each passage to monitor growth. On day 14 post-transduction, cell pellets were 1) collected for the viability arm, and 2) stained with CD47 antibody to sort for the top 50% and bottom 25% expression bins, described in detail below.

### Cell sorting for tiling screens

HT29 and/or A375 cells were transduced with virus for each of the Cas-containing vectors separately; 2 days after transduction, cells were selected with blasticidin for 14 days. Blasticidin was removed for one passage and cells were subsequently transduced with virus for the tiling guide library. 2 days after transduction, cells were selected with puromycin for 5 days. Following selection, cells were sorted on a SH800 Cell Sorter at varying time points. To prepare samples for sorting, cells were stained with a fluorophore-conjugated antibody targeting the respective cell surface marker gene, diluted 1:100 for 20-30 minutes on ice.

B2M: APC anti-human B2M antibody (Biolegend, 316312)

CD4: APC anti-human CD4 antibody (Biolegend, 357408)

CD26 (DPP4): FITC anti-human CD26 antibody (Biolegend, 302704)

CD46: FITC anti-human CD46 antibody (Biolegend, 315304)

CD47: FITC anti-human CD47 antibody (Biolegend, 323106)

CD55: FITC anti-human CD55 antibody (Biolegend, 311305)

CD59: FITC anti-human CD59 antibody (Biolegend, 304706)

CD63: PE anti-human CD63 antibody (Biolegend, 353004)

CD81: APC anti-human CD81 antibody (Biolegend, 349510)

CD97 (ADGRE5): FITC anti-human CD97 antibody (Biolegend, 336306)

CD274 (PD-L1): APC anti-human CD274 antibody (Biolegend, 329708)

LDLR: PE anti-human LDLR antibody (BD Biosciences, 565653)

Cells were washed with PBS two times to remove residual antibody and were resuspended in flow buffer (PBS, 2% FBS, 5 μM EDTA). Fluorophore signal was measured in the respective channel (APC-A, FITC-A, or PE-A). Gates were set such that the top ∼1% of cells were sorted into a high bin and the bottom ∼1% of cells were sorted into a low bin, into FBS. After sorting was completed, cells were centrifuged at 211 x g for 5 minutes. Following centrifugation, the FBS was aspirated and cells were resuspended in PBS.

### Genomic DNA isolation and sequencing

Genomic DNA (gDNA) was isolated using the KingFisher Flex Purification System with the Mag-Bind® Blood & Tissue DNA HDQ Kit (Omega Bio-Tek). For smaller cell pellets from sorted cell populations, gDNA was isolated using the NucleoSpin Blood Mini Kit (Macherey-Nagel). The gDNA concentrations were quantitated by Qubit.

For PCR amplification, gDNA was divided into 100 uL reactions such that each well had at most 10 ug of gDNA. Plasmid DNA (pDNA) was also included at a maximum of 100 pg per well. Per 96-well plate, a master mix consisted of 150 uL DNA Polymerase (Titanium Taq; Takara), 1 mL of 10x buffer, 800 uL of dNTPs (Takara), 50 uL of P5 stagger primer mix (stock at 100 uM concentration), 500 μL of DMSO (if used), and water to bring the final volume to 4 mL. Each well consisted of 50 uL gDNA and water, 40 uL PCR master mix, and 10 uL of a uniquely barcoded P7 primer (stock at 5 uM concentration). PCR cycling conditions were as follows: (1) 95°C for 1 minute; (2) 94°C for 30 seconds; (3) 52.5°C for 30 seconds; (4) 72°C for 30 seconds; (5) go to (2), x 27; (6) 72°C for 10 minutes. PCR primers were synthesized at Integrated DNA Technologies (IDT). PCR products were purified with Agencourt AMPure XP SPRI beads according to manufacturer’s instructions (Beckman Coulter, A63880), using a 1:1 ratio of beads to PCR product. Samples were sequenced on a HiSeq2500 HighOutput (Illumina) with a 5% spike-in of PhiX. For Cas12a guide constructs, a custom oligo was used for sequencing (oligo sequence: CTTGTGGAAAGGACGAAACACCGGTAATTTCTACTCTTGTAGAT).

### Flow cytometry assays for individual guide validation

HT29, HCT116, and/or A375 cells were transduced with virus for each of the Cas-containing vectors separately; 2 days after transduction, cells were selected with blasticidin for 14 days. Blasticidin was removed for one passage and cells were subsequently transduced with virus for guide-containing vectors. 2 days after transduction, cells were selected with puromycin for 5 days. Following selection, cells were visualized by flow cytometry on a CytoFLEX S Sampler at varying time points. To prepare samples for visualization, cells were stained and resuspended in flow buffer as described above. Fluorophore signal was measured in the respective channel (APC-A, FITC-A, or PE-A). Flow cytometry data were analyzed using FlowJo (v10.8.1). Gates were set such that ∼1% of cells score as APC-, FITC-, or PE-positive in the control condition (stained parental cells).

### *EMX1* sgRNA cloning

The selection of *EMX1* (target site: 5’ AAAGGTGAAAGAGAGATGGCT 3’; PAM sequence: 5’ GGGGGT 3’) was based on reported editing at the site by SaCas9, SlugCas9, and SauriCas9 in HEK293T cells (reported as target site “E4”)^33,34^. *EMX1*-targeting guide oligos for cloning into the all-in-one AAV constructs, described in Supplementary Data 2, were manufactured by GeneWiz and resuspended in water at a concentration of 100 uM. The two oligos were combined in equimolar amounts and phosphorylated using T4 PNK enzyme (New England Biolabs #M0201S) for 30 minutes at 37°C and then slowly annealed in a thermocycler to make the final double-stranded fragment with sticky ends for cloning into the all-in-one AAV constructs. The annealed oligos were diluted to a final concentration of 2 uM.

The all-in-one constructs (e.g., pRDA_917) were digested with BsmBI-v2 (New England Biolabs #R0739S) for two hours at 55°C. 1 uL of QuickCIP (New England Biolabs #M0525S) was added directly to the backbone digestions and incubated at 37°C for 30 minutes. The digested backbones were cleaned up using the Zymo DNA Clean & Concentrator Kit (Zymo Research #D4033) and quantified via Nanodrop. The final ligations with T4 DNA Ligase (New England Biosciences #M0202T) were set up using 25 ng of digested backbone plus 1 uL of the 2 uM annealed oligos in a final reaction volume of 10 uL. The ligation was allowed to proceed at room temperature for 1 hour, then 1 uL of each ligation reaction was transformed into 10 uL of NEB 5-alpha competent *E. coli* (New England Biosciences #C2987H) according to the manufacturer’s protocol. The transformed ligations were grown overnight at 37°C with ampicillin selection. Single colonies were used to inoculate ampicillin liquid cultures grown overnight with shaking at 37°C and plasmids were subsequently purified with the ZR Plasmid Miniprep kit (Zymo Research #D4016). The purified plasmids were screened by Sanger sequencing (GeneWiz) to ensure that the *EMX1*-targeting guide was correctly ligated using the following sequencing primer that partially anneals to the backbones: 5’ CTTCACCGAGGGCCTATTTC 3’.

### AAV production and purification

Maxipreps for the purpose of producing AAV packaging plasmids were purified from 200 mL overnight *E. coli* cultures using the EndoFree Plasmid Maxi Kit (Qiagen #12362). Low passage number (≤ passage 10) HEK293T cells were seeded in 15 cm cell culture dishes 18-24 hours before transfection and reached ∼80% confluency on the day of transfection. A single 15 cm dish was used to produce each individual virus at small scale. Each 15 cm dish was transfected with TransIT-LT1 transfection reagent (Mirus Bio #2300) according to the manufacturer’s protocol with the following plasmid amounts: 7 ug of the transfer plasmid (e.g., pRDA_917), 8 ug of the pAAV2/6 RepCap plasmid, and 10 ug of the pAdDeltaF6 helper plasmid.

48 hours after TransIT-LT1 transfection, the virus was harvested and purified. A cell scraper was used to remove cells from the bottom of the 15 cm dish and the entire cell-containing supernatant was collected in a conical tube. Chloroform (Millipore Sigma #650498) was added to the supernatant (10% of the initial volume). The supernatant-chloroform mix was shaken at 37°C for one hour, then NaCl was added to the solution to a final concentration of 1 M. The supernatant was then centrifuged at 20,000 x g for 15 minutes. All centrifuge steps took place at 4°C. The aqueous layer was carefully removed to a new tube and 10% weight/volume PEG8000 (Promega #V3011) was added and incubated on ice for 1 hour before centrifugation at 20,000 x g for 15 minutes. All supernatant was discarded and the pellet was resuspended in 5 mL of DPBS (Gibco #14190250). 5 uL of benzonase (Millipore Sigma #E1014-25KU) was added to the resuspended pellet along with 1 M MgCl2 to a final concentration of 1 mM. The benzonase digestion was incubated in a 37°C water bath for 30 minutes. After the incubation, an equal volume of chloroform was added, mixed, and centrifuged at 24,600 x g for 15 minutes. The aqueous layer was carefully collected and then buffer exchanged into DPBS and concentrated to ∼1 mL final volume using a 100 kDa molecular weight cutoff centrifugal filter unit (Millipore Sigma #UFC810024). The viral preps were aliquoted and stored at -80°C until use.

### AAV quantification by ddPCR

After purification, the viral titer of each prep was determined by droplet digital PCR (ddPCR) based on the method from Lock *et al.* 2014^64^. The BioRad QX200 system was used for droplet generation and droplet reading. Serial dilutions of the virus sample were prepared and used as template for the ddPCR reaction. Each ddPCR reaction was set up by adding the following: 10 uL of 2X ddPCR Supermix (no dUTP; BioRad #1863023), 2 uL of 10 uM ITR_Forward Primer, 2 uL of 10 uM ITR_Reverse Primer, 0.9 uL of 5 uM ITR_Probe, 2 uL of diluted virus prep, and 3.1 uL water, for a total volume of 20 uL. ddPCR primers and probes were as described in Supplementary Data 2.

After the reactions were set up, droplets were generated and either used immediately for the PCR step or stored at 4°C for up to 24 hours before the PCR step. PCR cycling conditions were as follows: (1) 95°C for 10 minutes; (2) 94°C for 30 seconds; (3) 58°C for 1 minute; (4) go to (2), x 35; (5) 98°C for 10 minutes; (6) 4°C hold.

After the PCR step, the BioRad QX200 was set up for absolute quantification of FAM-containing droplets. All ddPCR measurements for dilutions where the calculated number of copies per reaction did not exceed the number of droplets analyzed were used to calculate genome copy (GC)/mL concentration with the following formula and then averaged to determine the final titer for each prep: GC/mL = (ddPCR Copies per reaction * Dilution Factor * 1000)/volume of virus in reaction.

### AAV transductions

3.75e4 HEK293T cells were seeded in each well of 48-well plates. ∼24 hours after seeding, virus was added directly to each well to achieve the desired multiplicity of infection (MOI). Wells treated with DPBS (vehicle control) received a volume equal to the highest volume of virus used. 72 hours after addition of virus, the media was removed and genomic DNA was extracted using 30 uL/well of QuickExtract DNA Extraction Solution (Lucigen #QE09050) according to the manufacturer’s protocol. Briefly, the QuickExtract solution was added directly to cells after media removal. The solution was collected in PCR tubes and subjected to the thermocycler method specified by the manufacturer. The final volume of genomic DNA was brought to 100 uL after the thermocycler step and stored at -20°C until use.

MOIs for each viral prep were calculated based off of each well containing 7.5e5 cells (assuming a 24-hour doubling time). Specifically, the volume (mL) of virus prep needed to reach a certain MOI was calculated by multiplying the number of cells per well (7.5e5) by the desired MOI (50000, 100000, or 150000) then dividing this number by the calculated GC/mL concentration of the virus prep.

### *EMX1* editing analysis

Genomic DNA was used as the template DNA for a PCR with Q5 High-Fidelity 2X Master Mix (New England Biosciences #M0492L) to amplify the endogenous *EMX1* target site. *EMX1* primers were as described in Supplementary Data 2. Per well, each reaction consisted of 12.5 μL Q5 Master Mix, 10.5 uL water, 1 uL genomic DNA, 0.5 uL EMX1_Forward primer (stock at 25 uM), and 0.5 uL EMX1_Reverse primer (stock at 25 uM), for a total volume of 25 uL. PCR cycling conditions were as follows: (1) 98°C for 30 seconds; (2) 98°C for 10 seconds; (3) 65°C for 10 seconds; (4) 72°C for 15 seconds; (5) go to (2), x 34; (6) 72°C for 2 minutes; (7) 4°C hold. After completion of the thermocycler program, unpurified PCR products were sent for Sanger sequencing (GeneWiz) using the EMX1_Forward primer. The Sanger sequencing results were analyzed using Tracking of Indels by DEcomposition (TIDE)38 to calculate the amount of small indels (≤10 base pairs) in edited samples by comparison to an unedited control sample (PBS vehicle treated condition).

### *Drosophila* transgenics and crosses

Assembled UAS-DjCas13-2A-GFP test plasmid was transformed into Top10 chemically competent *E. coli*, miniprepped using the QIAprep Spin Miniprep Kit (Qiagen # 27104), eluted in injection buffer (100 µM NaPO4 and 5 mM KCl), and adjusted to 50 ng/µl. For phiC31-integration, plasmid was injected into *y v nos-phiC31-int; attP40* (for chromosome 2 insertions) or *y v nos-phiC31-int; attP2* (for chromosome 3 insertions). Injected male G0 flies were crossed with *y w; Gla/CyO or y w; Dr e/TM3, Sb* to identify transformants (marked by *w+*) and remove the integrase from the X chromosome, and subsequently balanced.

Balanced *UAS-DjCas13-2A-GFP* transformants were crossed to the wing-specific *nubbin-GAL4* driver line (Perrimon Lab). 3^rd^ instar wing discs from *nubbin-Gal4; UAS-DjCas13-2A-GFP* or *nubbin-Gal4* control larvae were dissected in PBS, fixed in 4% formaldehyde, rinsed in PBS and mounted on glass slides with vectashield (H-1000; Vector Laboratories) under a coverslip. Images of mounted wing discs and fly eyes were acquired with a Zeiss Axio Zoom V16 fluorescence microscope.

### *Drosophila* cell culture

*Drosophila* S2R+ cells^41^ or S2R+ cells with *mCherry* and *attP* integrated into the *clic* locus (aka PT5 cells)^42^, were cultured at 25°C, using Schneider’s medium (21720-024; Thermo Fisher Scientific) with 10% fetal bovine serum (A3912; Sigma [Sigma Chemical], St. Louis, MO) and 50 U/ml penicillin/streptomycin (15070-063; Thermo Fisher Scientific). Plasmid transfections were performed in six-well tissue-culture treated dishes at 1.8 × 10^6 cells/ml using Effectene (301427; QIAGEN) following the manufacturer’s instructions. For antibiotic selection, media was replaced with antibiotic-containing media 24 hours after transfection. Five days after transfection, cells were harvested and diluted 1:7.5 into a T75 flask containing antibiotic-containing media. After seven days, cells were centrifuged at 250g for 10min, resuspended in 2ml antibiotic-containing media, and serially diluted into a six-well dish. Plates were monitored every few days until wells contained ∼10-100 robustly growing colonies of selected cells and were confluent. Selected cells were resuspended and expanded into T25 flasks. Cells were imaged using an EVOS M5000 Microscope Imaging System (AMF5000). GFP was imaged using EVOS™ Light Cube, GFP 2.0 (Invitrogen, AMEP4951), mCherry was imaged using EVOS™ Light Cube, RFP 2.0 (Invitrogen, AMEP4952), and TagRFP657 fluorescence was detected using EVOS™ Light Cube, Texas Red 2.0 (Invitrogen, AMEP4955).

For experiments involving assembled plasmid *Act5C-BE-SpRY-2A-TagRFP657*, 2µg of plasmid was transfected into S2R+ cells in a single well of a six-well plate. 200µg/ml Hygromycin (EMD Millipore 400051-1MU) was used to select for stable cell lines by random genome integration.

For experiments involving assembled plasmid Act5C-*ZIM3-dCas9-2A-puro-2A-GFP*, PT5 cells were transfected with a plasmid mix containing *ZIM3-dCas9-2A-puro-2A-GFP* (200ng) and PhiC31 integrase plasmid (*pBS130*; Addgene, 26290) (200ng), or *ZIM3-dCas9-2A-puro-2A-GFP* (200ng) and *pCFD3* (Addgene, 49410) (200ng) (no-integrase control transfection). 5µg/ml Puromycin (EMD Millipore 540411-100MG) was used to select for stable cell lines by PhiC31 integration into the *clic attP* site.

## QUANTIFICATION AND STATISTICAL ANALYSIS

### Screen analysis

Guide sequences were extracted from sequencing reads by running PoolQ (https://portals.broadinstitute.org/gpp/public/software/poolq). Reads were counted by alignment to a reference file of all possible guide RNAs present in the library. The read was then assigned to a condition (e.g., a well on the PCR plate) on the basis of the 8 nucleotide index included in the P7 primer. Following deconvolution, the resulting matrix of read counts was first normalized to reads per million within each condition by the following formula: read per guide RNA / total reads per condition x 1e6. Reads per million was then log2-transformed by first adding one to all values, which is necessary in order to take the log of guides with zero reads. Prior to further analysis, we filtered out guides for which the log-normalized reads per million of the pDNA was >3 standard deviations from the mean. For the high-throughput GG-assembled screens, barcode sequences were used instead of guide RNAs for all analyses.

We then calculated the log2-fold-change (LFC) between conditions. All dropout conditions were compared to early time point (ETP) samples; for flow-sorted screens, low-expressing populations were compared to high-expressing populations. We then assessed the correlation between LFC values of replicates. Delta-log2-fold-change values (dLFCs) were calculated by subtracting intergenic LFCs from targeting LFCs.

### Data visualization

Figures were created with Python3, FlowJo 10.9.0, and GraphPad Prism (version 10). Schematics were created with BioRender.

### Statistical analysis

All correlation coefficients were calculated in Python.

## DATA AVAILABILITY

DepMap 22Q4 was used for all analyses. The read counts for screening data are provided as Supplementary Data.

## CODE AVAILABILITY

PoolQ, used to deconvolute sequencing reads, is available to download from the GPP portal.

## SUPPLEMENTAL DATA

**Title:** Supplementary Data 1.

**Description:** Guide sequences - positive control guide sequences from flow cytometry-based tiling screens. Associated with Figure 2.

**Title:** Supplementary Data 2.

**Description:** Oligo and primer sequences. Associated with Figure 3.

**Title:** Supplementary Data 3.

**Description:** Pooled GG screens - read counts, library annotation, replicate correlations. Associated with Figure 4.

### Supplementary Note 1: Creation of pF vectors

All pF plasmids were made by gene synthesis into the EcoRV site of pUC57-Kan (or a variant thereof that also lacked BsmBI sites). The starting plasmid sequence is provided below.

To create a new pF vector, it will likely be helpful to first download an existing pF vector from the Fragmid website that uses the same module set as the desired new vector. For example, to make a new Cas protein, download pF_AA007 (Cas9 from *S. pyogenes*) and replace the Cas9 coding sequence with that of the new protein. Remove any internal BbsI or BsmBI sites with silent mutations.

Be sure to retain the module sets, which are labeled in the .gb files as GG-#F and GG-#R. Note that some module sets include extra nucleotides to complete the coding frame of the Gly-Ser linker between translated fragments. These should be included in the synthesis order.

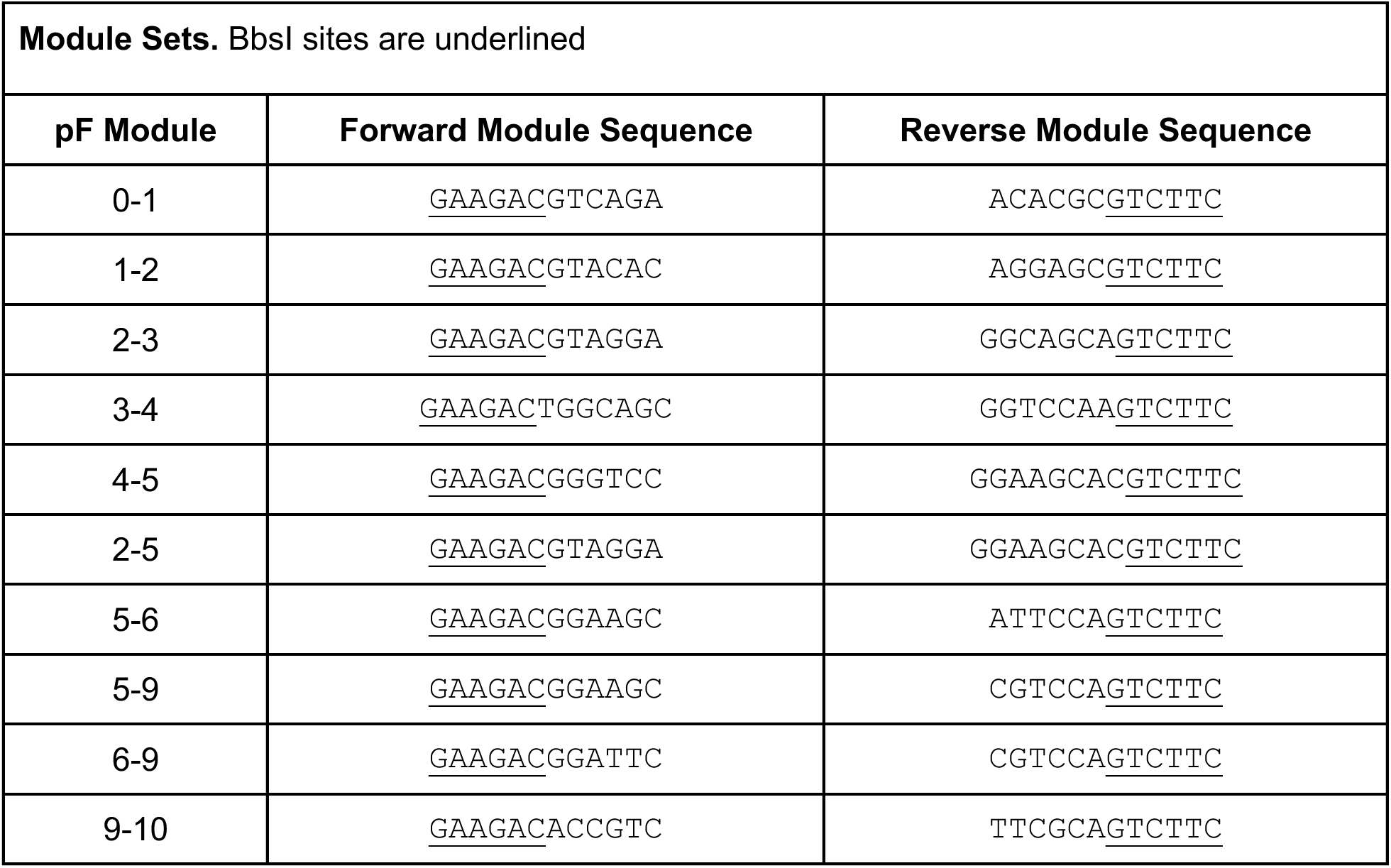

### Schematic

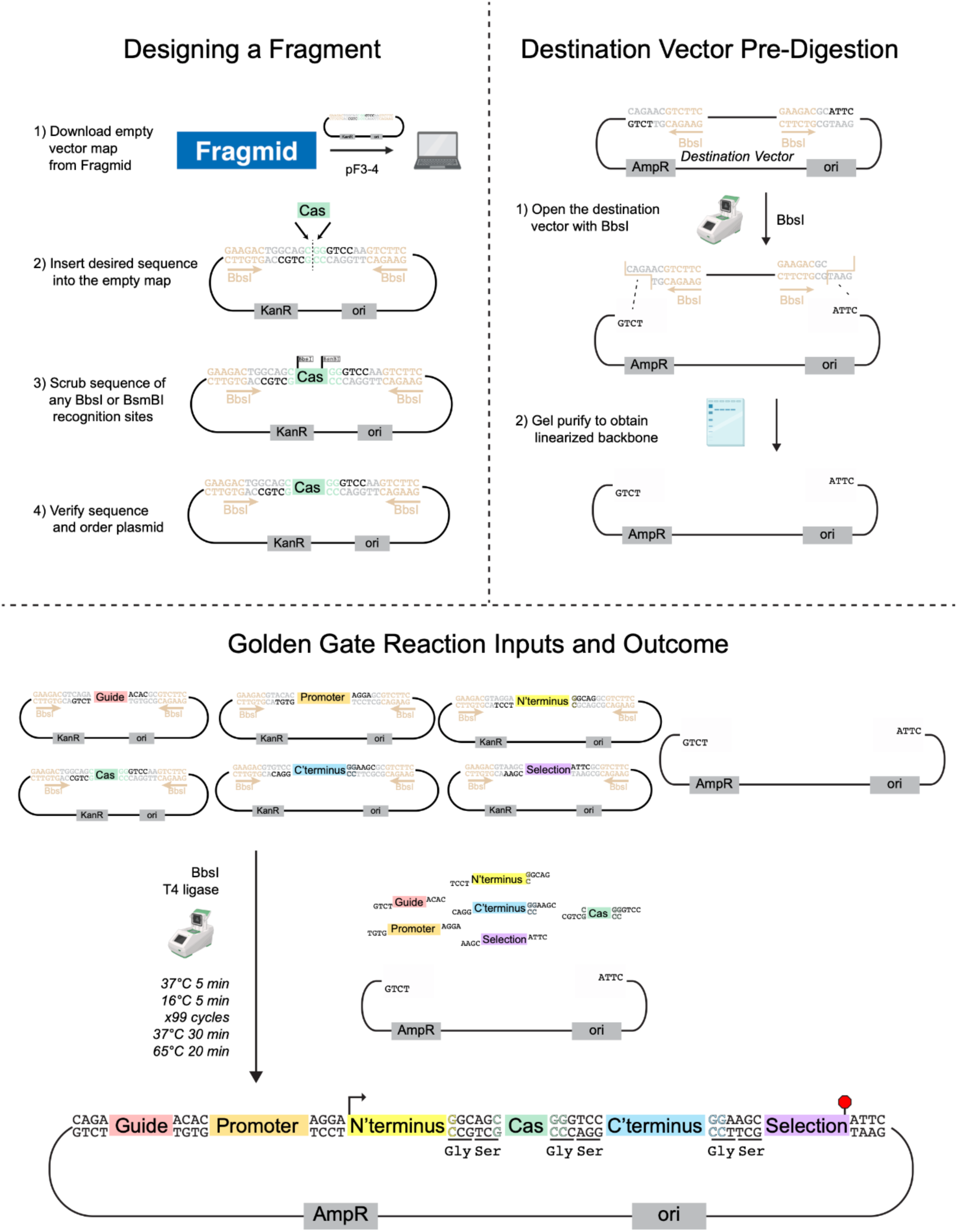

### Vector sequences

#### pUC57-Kan

tcgcgcgtttcggtgatgacggtgaaaacctctgacacatgcagctcccggagacggtcacagcttgtctgtaagcggatgccgggagcagacaagcccgtcagggcgcgtcagcgggtgttggcgggtgtcggggctggcttaactatgcggcatcagagcagattgtactgagagtgcaccatatgcggtgtgaaataccgcacagatgcgtaaggagaaaataccgcatcaggcgccattcgccattcaggctgcgcaactgttgggaagggcgatcggtgcgggcctcttcgctattacgccagctggcgaaagggggatgtgctgcaaggcgattaagttgg gtaacgccagggttttcccagtcacgacgttgtaaaacgacggccagagaattcgagctcggtacctcgcgaatacatcta**gatatc** ggatcccgggcccgtcgactgcagaggcctgcatgcaagcttggtgtaatcatggtcatagctgtttcctgtgtgaaattgttatccgctcacaattccacacaacatacgagccggaagcataaagtgtaaagcctggggtgcctaatgagtgagctaactcacattaattgcgttgcgctcactgcccgctttccagtcgggaaacctgtcgtgccagctgcattaatgaatcggccaacgcgcggggagaggcggtttgcgtattgggcgctcttccgcttcctcgctcactgactcgctgcgctcggtcgttcggctgcggcgagcggtatcagctcactcaaaggcggtaatacggttatccacagaatcaggggataacgcaggaaagaacatgtgagcaaaaggccagcaaaaggccaggaaccgtaaaaaggccgcgttgctggcgtttttccataggctccgcccccctgacgagcatcacaaaaatcgacgctcaagtcagaggtggcgaaacccgacaggactataaagataccaggcgtttccccctggaagctccctcgtgcgctctcctgttccgaccctgccgcttaccggatacctgtccgcctttctcccttcgggaagcgtggcgctttctcatagctcacgctgtaggtatctcagttcggtgtaggtcgttcgctccaagctgggctgtgtgcacgaaccccccgttcagcccgaccgctgcgccttatccggtaactatcgtcttgagtccaacccggtaagacacgacttatcgccactggcagcagccactggtaacaggattagcagagcgaggtatgtaggcggtgctacagagttcttgaagtggtggcctaactacggctacactagaagaacagtatttggtatctgcgctctgctgaagccagttaccttcggaaaaagagttggtagctcttgatccggcaaacaaaccaccgctggtagcggtggtttttttgtttgcaagcagcagattacgcgcagaaaaaaaggatctcaagaagatcctttgatcttttctacggggtctgacgctcagtggaacgaaaactcacgttaagggattttggtcatgagattatcaaaaaggatcttcacctagatccttttaaattaaaaatgaagttttaaatcaagcccaatctgaataatgttacaaccaattaaccaattctgattagaaaaactcatcgagcatcaaatgaaactgcaatttattcatatcaggattatcaataccatatttttgaaaaagccgtttctgtaatgaaggagaaaactcaccgaggcagttccataggatggcaagatcctggtatcggtctgcgattccgactcgtccaacatcaatacaacctattaatttcccctcgtcaaaaataaggttatcaagtgagaaatcaccatgagtgacgactgaatccggtgagaatggcaaaagtttatgcatttctttccagacttgttcaacaggccagccattacgctcgtcatcaaaatcactcgcatcaaccaaaccgttattcattcgtgattgcgcctgagcgagacgaaatacgcgatcgctgttaaaaggacaattacaaacaggaatcgaatgcaaccggcgcaggaacactgccagcgcatcaacaatattttcacctgaatcaggatattcttctaatacctggaatgctgtttttccggggatcgcagtggtgagtaaccatgcatcatcaggagtacggataaaatgcttgatggtcggaagaggcataaattccgtcagccagtttagtctgaccatctcatctgtaacatcattggcaacgctacctttgccatgtttcagaaacaactctggcgcatcgggcttcccatacaagcgatagattgtcgcacctgattgcccgacattatcgcgagcccatttatacccatataaatcagcatccatgttggaatttaatcgcggcctcgacgtttcccgttgaatatggctcataacaccccttgtattactgtttatgtaagcagacagttttattgttcatgatgatatatttttatcttgtgcaatgtaacatcagagattttgagacacgggccagagctgca

#### pUC57-Kan_BsmBI-free

tcgcgcgtttcggtgatgacggtgaaaacctctgacacatgcagctcccgcaggcggtcacagcttgtctgtaagcggatgccgggagcagacaagcccgtcagggcgcgtcagcgggtgttggcgggtgtcggggctggcttaactatgcggcatcagagcagattgtactgagagtgcaccatatgcggtgtgaaataccgcacagatgcgtaaggagaaaataccgcatcaggcgccattcgccattcaggctgcgcaactgttgggaagggcgatcggtgcgggcctcttcgctattacgccagctggcgaaagggggatgtgctgcaaggcgattaagttgg gtaacgccagggttttcccagtcacgacgttgtaaaacgacggccagagaattcgagctcggtacctcgcgaatacatcta**gatatc** ggatcccgggcccgtcgactgcagaggcctgcatgcaagcttggtgtaatcatggtcatagctgtttcctgtgtgaaattgttatccgctcacaattccacacaacatacgagccggaagcataaagtgtaaagcctggggtgcctaatgagtgagctaactcacattaattgcgttgcgctcactgcccgctttccagtcgggaaacctgtcgtgccagctgcattaatgaatcggccaacgcgcggggagaggcggtttgcgtattgggcgctcttccgcttcctcgctcactgactcgctgcgctcggtcgttcggctgcggcgagcggtatcagctcactcaaaggcggtaata cggttatccacagaatcaggggataacgcaggaaagaacatgtgagcaaaaggccagcaaaaggccaggaaccgtaaaaaggccgcgttgctggcgtttttccataggctccgcccccctgacgagcatcacaaaaatcgacgctcaagtcagaggtggcgaaacccgacaggactataaagataccaggcgtttccccctggaagctccctcgtgcgctctcctgttccgaccctgccgcttaccggatacctgtccgcctttctcccttcgggaagcgtggcgctttctcatagctcacgctgtaggtatctcagttcggtgtaggtcgttcgctccaagctgggctgtgtgcacgaaccccccgttcagcccgaccgctgcgccttatccggtaactatcgtcttgagtccaacccggtaagacacgacttatcgccactggcagcagccactggtaacaggattagcagagcgaggtatgtaggcggtgctacagagttcttgaagtggtggcctaactacggctacactagaagaacagtatttggtatctgcgctctgctgaagccagttaccttcggaaaaagagttggtagctcttgatccggcaaacaaaccaccgctggtagcggtggtttttttgtttgcaagcagcagattacgcgcagaaaaaaaggatctcaagaagatcctttgatcttttctacggggtctgacgctcagtggaacgaaaactcacgttaagggattttggtcatgagattatcaaaaaggatcttcacctagatccttttaaattaaaaatgaagttttaaatcaagcccaatctgaataatgttacaaccaattaaccaattctgattagaaaaactcatcgagcatcaaatgaaactgcaatttattcatatcaggattatcaataccatatttttgaaaaagccgtttctgtaatgaaggagaaaactcaccgaggcagttccataggatggcaagatcctggtatcggtctgcgattccgactcgtccaacatcaatacaacctattaatttcccctcgtcaaaaataaggttatcaagtgagaaatcaccatgagtgacgactgaatccggtgagaatggcaaaagtttatgcatttctttccagacttgttcaacaggccagccattacgctcgtcatcaaaatcactcgcatcaaccaaaccgttattcattcgtgattgcgcctgagccaggcgaaatacgcgatcgctgttaaaaggacaattacaaacaggaatcgaatgcaaccggcgcaggaacactgccagcgcatcaacaatattttcacctgaatcaggatattcttctaatacctggaatgctgtttttccggggatcgcagtggtgagtaaccatgcatcatcaggagtacggataaaatgcttgatggtcggaagaggcataaattccgtcagccagtttagtctgaccatctcatctgtaacatcattggcaacgctacctttgccatgtttcagaaacaactctggcgcatcgggcttcccatacaagcgatagattgtcgcacctgattgcccgacattatcgcgagcccatttatacccatataaatcagcatccatgttggaatttaatcgcggcctcgacgtttcccgttgaatatggctcataacaccccttgtattactgtttatgtaagcagacagttttattgttcatgatgatatatttttatcttgtgcaatgtaacatcagagattttgagacacgggccagagctgca

---

**Supplementary Figure 1.**
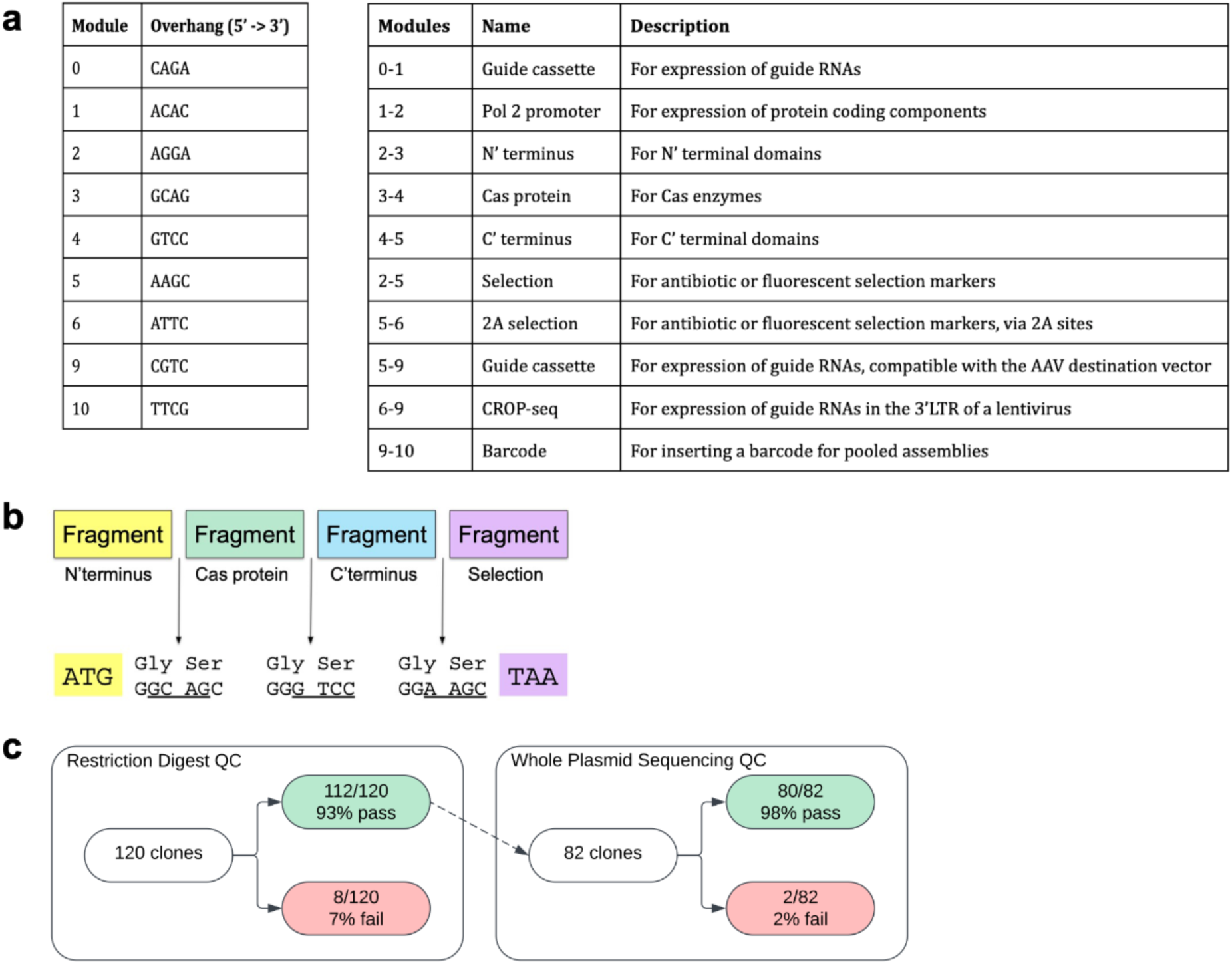
Modular vector overview. **A**) Table of BbsI overhang sequences and associated module numbers (left). Table of fragment types, descriptions, and module numbers (right). **B**) Schematic depicting glycine-serine linkers encoded by BbsI overhang sequences for coding fragment types. **C**) Schematic depicting GG assembly fidelity at multiple QC checkpoints.

**Supplementary Figure 2.**
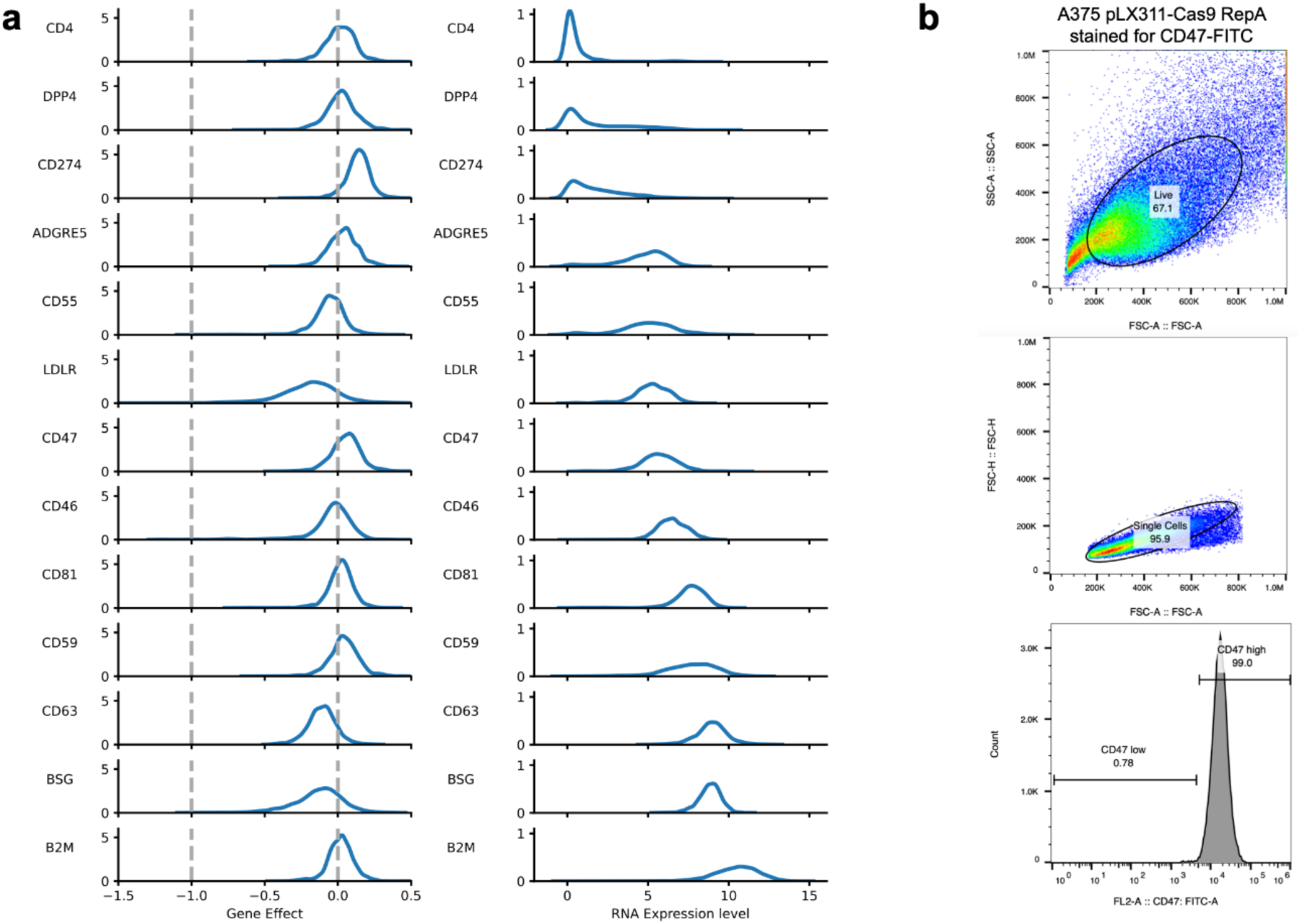
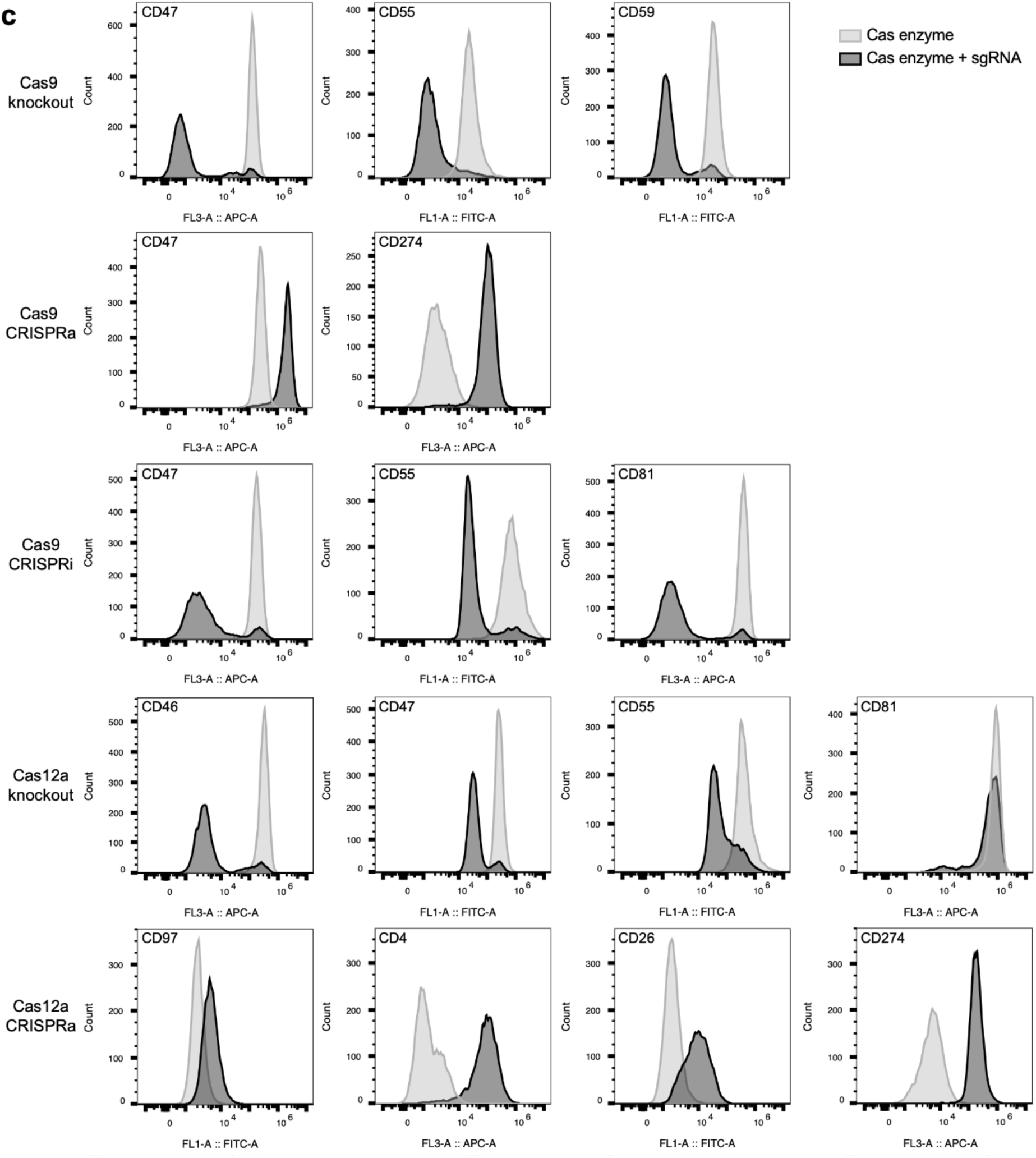
Tiling screens for identification of positive control guides. **A**) Gene effect scores (left) and RNA expression levels (right) from the DepMap for the 13 cell surface marker genes targeted by tiling libraries. Dashed gray line at gene effect score of -1 corresponds to the median effect of common essential genes; dashed gray line at gene effect score of 0 corresponds to no viability effect. **B**) Representative gating strategy used to assess fluorescence in the FITC channel. Stained cells were gated first for live cells (top). This live cell population was then gated to exclude doublets (middle), and the single cell population was then sorted on FITC (CD47) fluorescence (bottom). **C**) Histograms showing cell surface marker expression levels (APC or FITC channel) when targeted in validation experiments for each enzyme-technology combination. For the EnAsCas12a data, all four guides are expressed on the same array. Data from one representative replicate shown for each guide.

**Supplementary Figure 3.**
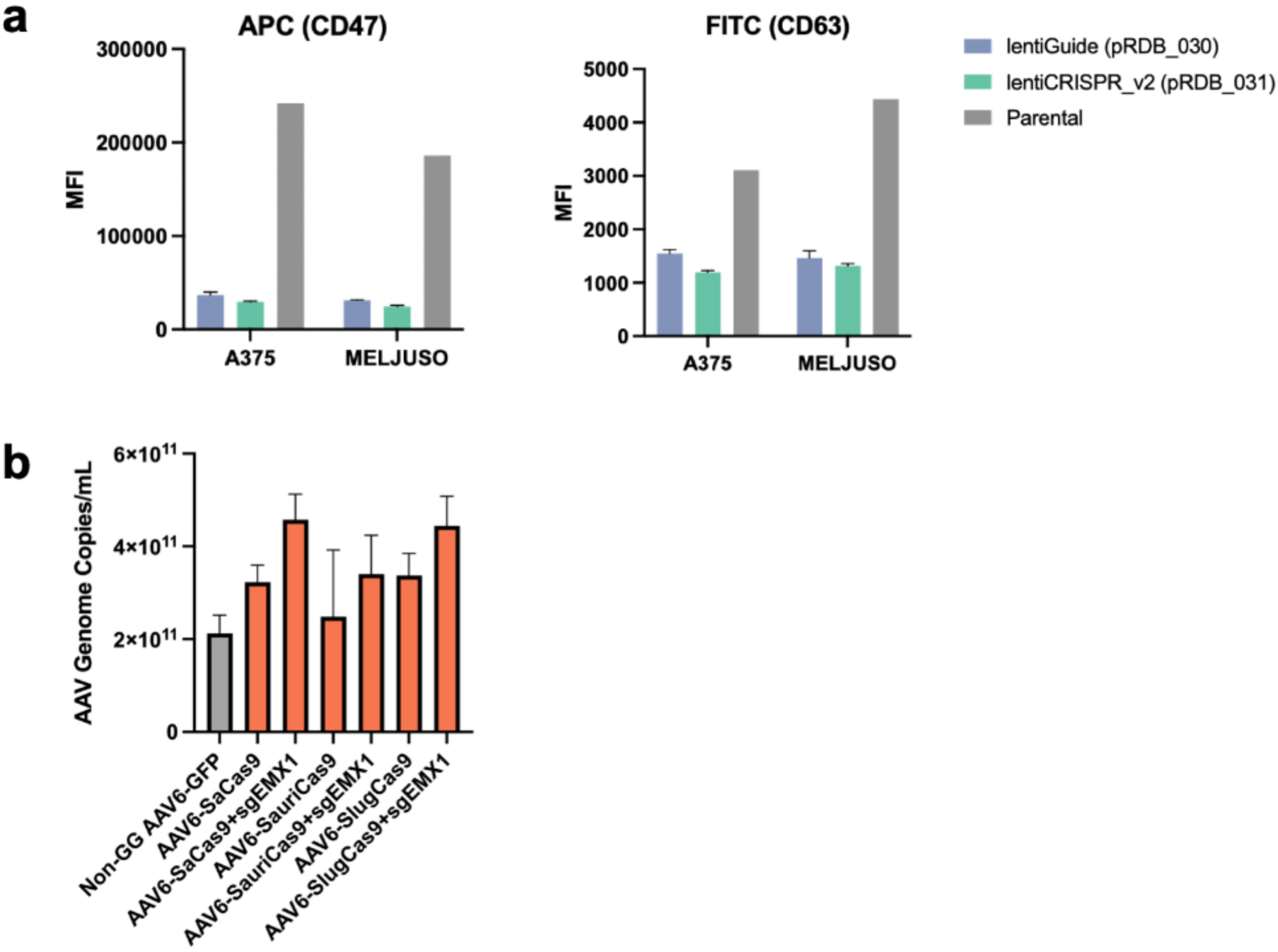
Applications for viral delivery. **A**) Comparison of CD47 and CD63 knockout activity in A375 and MelJuSo cells expressing an all-in-one EnAsCas12a construct assembled in the lentiGuide and lentiCRISPR_v2 destination vectors. Barplots show CD47 MFI values (left) and CD63 MFI values (right). Stained parental cells are depicted in gray. **B**) Barplot depicting AAV viral titer (genome copies/mL) quantified by ddPCR for SaCas9, SauriCas9, and SlugCas9 constructs, with and without the 21 nt *EMX1* guide cloned into the AAV destination vector. A GFP control construct cloned into a non-GG destination vector is depicted in gray.

**Supplementary Figure 4.**
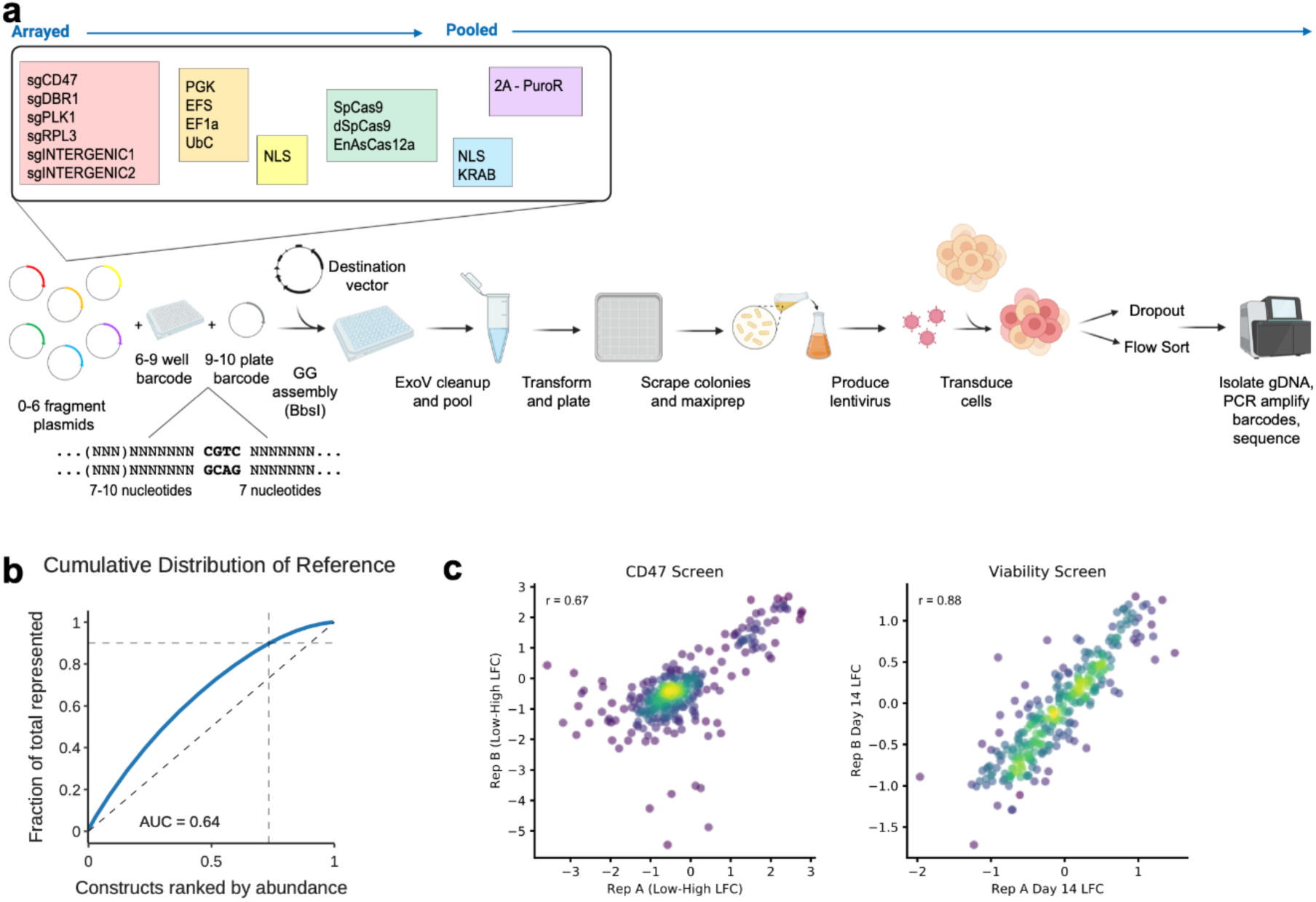
Pooled Golden Gate assembly. **A**) Schematic of partially-pooled GG assembly, including fragments used and barcoding strategy. B) Cumulative distribution of barcodes in pooled GG library pDNA. C) Replicate correlations (Pearson’s r) for the pooled GG library screened in MelJuSo cells in duplicate for the CD47-sort screen (left) and viability screen (right).

**Supplementary Figure 5.**
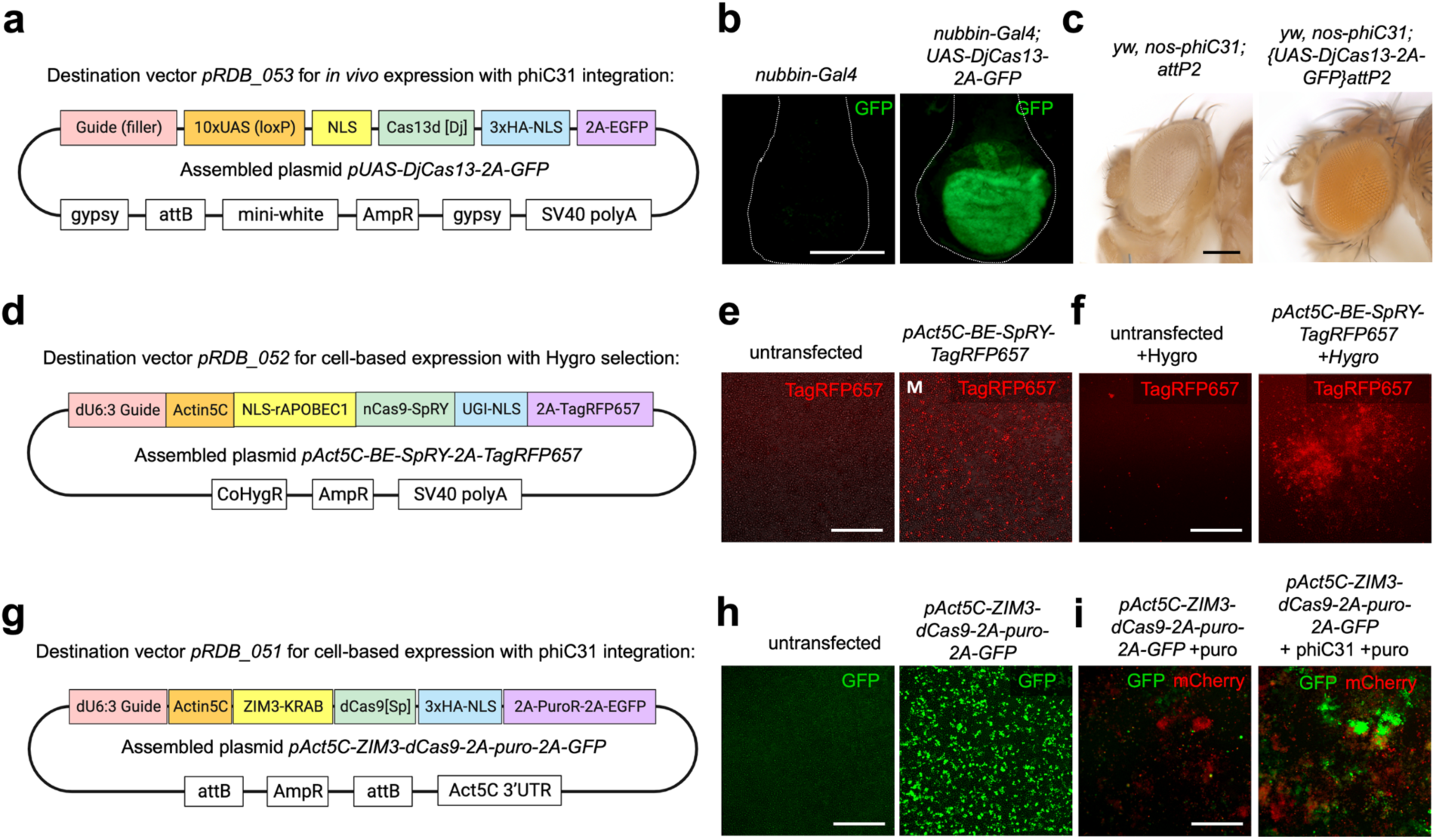
Applications for *Drosophila* cell-based and in vivo expression. **A**) Schematic of *pUAS-DjCas13-2A-GFP*, assembled in destination vector *pRDB_053*, for *in vivo* expression. **B**) Wing imaginal discs from *nubbin-Gal4*; *UAS-DjCas13-2A-GFP*, but not *nubbin-Gal4* controls, express GFP in the wing pouch. C) Injection of *pUAS-DjCas13-2A-GFP into yw, nos-phiC31*; *attP2* flies produces transformants with orange eyes. **D**) Schematic of *pAct5C-BE-SpRY-2A-TagRFP657* assembled in destination vector *pRDB_052*, for cell-based expression with Hygromycin selection. **E**) S2R+ cells transfected with *pAct5C-BE-SpRY-2A-TagRFP657* express TagRFP657 (red). **F**) Hygromycin selects for S2R+ cells expressing *pAct5C-BE-SpRY-2A-TagRFP657* (red). **G**) Schematic of *pAct5C-ZIM3-dCas9-2A-puro-2A-GFP* assembled in destination vector *pRDB_051*, for cell-based expression with phiC31 integration. **H**) S2R+ cells transfected with *pAct5C-ZIM3-dCas9-2A-puro-2A-GFP* express GFP (green). I) S2R+ PT5 cells transfected with *pAct5C-ZIM3-dCas9-2A-puro-2A-GFP* + phiC31, integrate the plasmid into the *attP* sites, (marked by increased GFP, and reduced mCherry fluorescence). Scale bars, 100mm (B, C); 300mm (E, F, H, I).

